# A method for analysing tissue motion and deformation during mammalian organogenesis

**DOI:** 10.1101/2025.06.27.661906

**Authors:** Morena Raiola, Isaac Esteban, Kenzo Ivanovitch, Miquel Sendra, Miguel Torres

**Author notes:** Institute for Biomedical Engineering, ETH Zürich, Zürich 8093, Switzerland. Developmental Biology of Birth Defects, Institute of Child Health, University College London, 30 Guilford Street, London WC1N 1EH, UK. CNRS, UTLN, LIS 7020, Turing Centre for Living Systems, Aix Marseille University, Marseille, France.

## Abstract

Understanding tissue morphogenesis is an important goal in developmental biology and tissue engineering. Accurately describing tissue deformation processes and how cell rearrangements contribute to these is a challenging task. Live analysis of morphogenesis in 3D is frequently used to obtain source data that allow to extract such features from developing organs. However, several limitations are encountered when applying these methodologies to mammalian embryos. The mouse embryo is the most frequently used model, but most studies use a very limited number of specimens and present only individual acquisitions due to constraints imposed by embryo culture and imaging. Here we used data from live analysis of embryonic heart development to develop methodologies that allow to 1) deduce tissue deformation and cell movements from image intensity flows, 2) stage and register data from several specimens/acquisitions to a dynamic consensus model of heart development, 3) statistically calculate and map to the model tissue growth and anisotropy and 4) generate a digital model of heart development that allows *in-silico* pseudo-cell fate mapping. Our methodology will greatly facilitate the understanding of morphogenetic processes underlying mammalian organogenesis.

## INTRODUCTION

Recent advances in microscopy physics and the development of fluorescent proteins detectable in living cells have revolutionized *in vivo* observation of organogenesis in mammalian embryos (Dominguez et al., 2023; Gómez et al., 2023; Stower & Srinivas, 2021; Zhu et al., 2020; Ivanovitch et al., 2017. This imaging process generates high-resolution spatio-temporal 3D+t data, providing a digital representation of organ morphogenesis at tissue and cellular scale. However, extracting information from these data necessitates *ad-hoc* image processing and data analysis techniques.

Characterizing the movement and deformation patterns of developing tissues poses a significant challenge in developmental biology. This complexity arises from the variability of organ morphogenesis and the inherent limitations of current imaging approaches (Raiola et al., 2023). Live imaging often falls short in capturing the full timeline of organ development or the entire region of interest within a single acquisition. To balance spatial and temporal resolution with field size in 3D+t imaging and enable cell tracking, data acquisition is typically limited to a few hours to avoid signal degradation and embryo damage. As a result, several acquisitions that cover different developmental stages have to be integrated to study the full developmental timeline (Ivanovitch et al., 2017).

Deep tissue imaging is further constrained by limited laser penetration in non-transparent specimens, such as mouse embryos, resulting in incomplete sampling. In addition, embryo movement during growth can also cause sample to drift out of focus, leading to the loss of peripheral regions of the organ of interest during. Here, we developed a new methodology to cope with these challenges using early mouse heart development as a model. Although heart morphogenesis is a generally consistent and robust process that transforms the early heart primordium into a functional four-chambered organ it exhibits significant in shape and maturation rates among individual embryos (Esteban et al., 2022, Ivanovitch et al., 2017). To overcome these challenges, we developed a systematic approach to quantify and compare myocardial motion and tissue deformation. Our method integrates live imaging and high-resolution static confocal images to address limitations of temporal resolution and variability in tissue morphology. The framework enables non-invasive analysis of tissue motion and deformation directly from individual live imaging datasets.

To synchronize the motion and deformation data across specimens, we developed a staging system that maps temporally and spatially 3D live images to a previously described pseudodynamic Atlas of tissue geometry during heart morphogenesis (Esteban et al., 2022). To account for variability in heart shape and enable comparisons of deformation patterns, we implemented a registration process that projects each individual specimen to the synchronous Atlas geometry, which serves as a consistent spatial reference.

By integrating live images into a unified spatiotemporal framework, we reconstructed cumulative tissue deformation and generated the first *in-silico* fate map of early heart morphogenesis. This allowed us to propose a new model for mouse heart tube morphogenesis. Our methodology provides a robust framework for analyzing organ morphogenesis from live imaging, offering tools to describe the cellular bases of tissue deformation, map spatial patterns, and quantify variability in developmental processes.

## RESULTS

We have developed a workflow that estimates motion profiles at individual cell level and combines multiple live images in time and space to quantify the deformation patterns and their variability during early mammalian heart development. Our workflow, illustrated in Figure 1, consists of four key steps: (1) Estimating individual live image motion to describe heart tube shaping as a continuous process (blue shapes); (2) Integrating multiple live images into a consensus spatiotemporal reference by aligning each continuous motion profile with a published 3D+t Atlas (Esteban et al., 2022) (red shapes); (3) Quantifying tissue deformation during early morphogenesis (yellow shapes); and (4) Creating an *in-silico* fate map to analyse heart tube morphogenesis at both pseudo-cellular and regional levels (green shapes).

**Figure 1:**
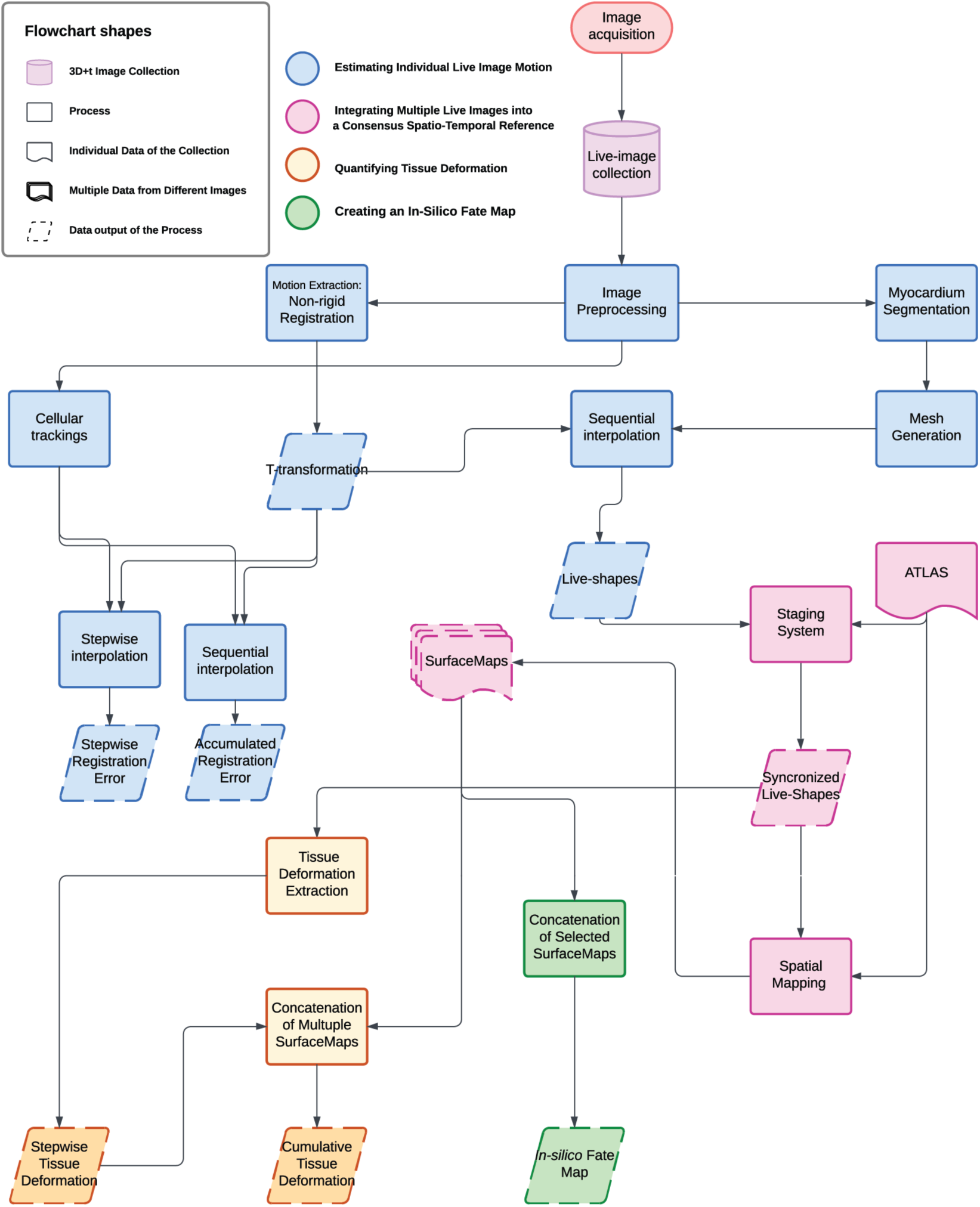
Workflow to Extract and Define Myocardial Motion and Deformation Patterns During Early Heart Morphogenesis. A comprehensive pipeline for the 3D imaging of early heart morphogenesis. This figure provides an overview of the workflow used in this study. For full details of each methodological step, please refer to the STAR Methods section. The computational workflow comprises four main components: 1) Estimating Individual Live Image Motion (blue shapes): Following image preprocessing, we extracted the underlying motion using a non-rigid registration algorithm, resulting in a set of T-transformation. The accuracy of motion detection was evaluated by comparing tracked cells to computed tracking using a stepwise and sequential interpolation. The myocardium tissue was segmented at one time point, and a mesh was generated. By interpolating this mesh with the set of T-transformation, we derived Live-Shape, a continuous description of heart tissue motion. 2) Integrating Multiple Live Images into the Atlas (red shapes): Individual live image motions were integrated into a high-resolution Atlas. This involved a staging system to synchronize Live-Shape collections along the Atlas time reference and a spatial mapping strategy to project staged Live-Shapes into the Atlas spatial framework. The result was a set of SurfaceMap, representing the motion of each specimen within the Atlas. 3) Quantifying Tissue Deformation (yellow shapes): Individual tissue deformation patterns were extracted, mapped into the Atlas, and variability was assessed. We defined stepwise tissue deformation and cumulative deformation to quantify these changes over time. 4) Creating an In-Silico Fate Map (green shapes): An *in-silico* fate map of the myocardium was constructed for the developmental window between E7.75 and E8.25 by concatenating motion profiles, providing insights into the spatial and temporal dynamics of early heart morphogenesis. Our dataset includes multiple specimens raging from E7.75 to E8.25 (12 hours). Two different Rosa26Rtdtomato^+/-^, Nkx2.5eGFP specimens are aligned on a pseudo-timeline, representing the transition from the cardiac crescent to the linear heart tube.

### Estimating Individual Live Image Motion

We used both published (Ivanovitch et al., 2017) and new datasets consisting of 3D+t live images. The dataset includes 16 embryos at different stages of heart development, from the cardiac crescent to the linear heart tube (Table S1). During this period the heart tissues undergo complex reorganization with a change in topology that converts a flat epithelial monolayer into a tube with two inflow (IFT) and outflow (OTF) tracts.

The imaged embryos were categorized based on their genetic reporters: some carried the Nkx2.5-GFP allele (Wu & Sato, 2008), while others had a combination of the Nkx2.5Cre allele (Stanley et al., 2004) and *Rosa26* fluorescent reporter alleles (Table S1). To facilitate cell tracking, Nkx2.5-GFP carriers also had cells labelled at random through the recombination of the Rosa26R Cre-reporter allele, induced by tamoxifen activation of CreERT2 from the RERT allele. Additional embryos were labelled with Mesp1Cre to report all mesodermal cells or Islet1Cre to mark the second heart field.

The images were stored as hyperstacks in the format I(x, y, z, c, t), with an xy resolution of 1024x1024 pixels and extending in the z-dimension to encompass the entire cardiogenic domain. Each frame had an xy resolution of 0.593 μm x 0.593 μm, and the axial displacement along the z-axis ranged from 3 to 6 μm. The duration of the acquisitions varied between 2 and over 10 hours, with frame rates ranging from 4 to 24 minutes depending on the specimen.

We estimated the motion of the heart from live images using a non-rigid registration technique. Specifically, we adapted the MIRT algorithm, (Myronenko et al., 2007) to pre-processed 3D+t images (see STAR Methods-Image Preprocessing, MIRT algorithm). We implemented this intensity-based approach sequentially, where, at each iteration, the algorithm aligns pairs of images frame-by-frame and extracts the underlying T-transformations that reproduce the flow of the image 3D intensity map (Figure 2A). This sequential implementation returns a set of transformation fields {Ti}, enabling the tracking of the positions of any point in an image throughout the video.

**Figure 2:**
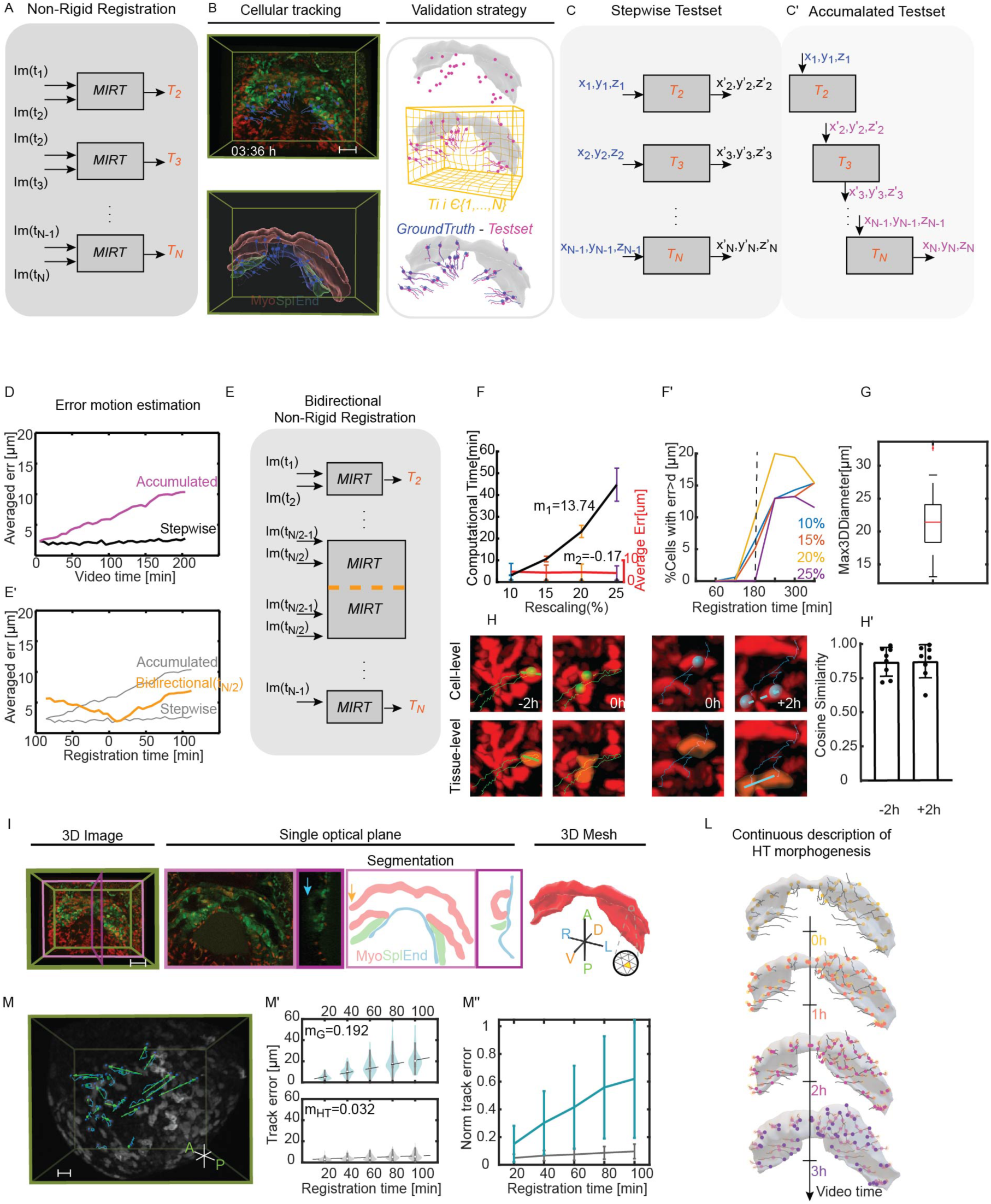
Estimating individual live image motion to describe heart tube shaping as a continuous process. (A), MIRT algorithm was applied frame-by-frame (I(t_i_)-I(t_i+1_)), extracting the set of deformation fields (T_1_, T_2_, …, T_N_) underlying the live images. (B), The top image illustrates the automatic cell tracking (blue tracks) on a live image (Nkx2.5eGFP; Rosa26Rtdtomato^+/-^). The bottom image is an Imaris reconstruction of the HT and the relative automatic cell tracking. This tracking serves as the ground truth dataset. On the left, there is a representation of the validation strategy. By applying the deformation fields, we built our test set (pink lines). We computed the Eulerian distances between the ground truth and the test set to assess the accuracy of the registration method. The scale bar is 100 µm and the time is represented in [hh:mm]. (C), The graph illustrates the method for generating the stepwise test set. For each centroid in the ground truth dataset, the T-transformation determines its new position (x′, y′, z′) at the next time point. (C’), The sequential test set is generated by sequentially interpolating the initial cell positions (x, y, z, t_0_) (shown in blue) using the transformation set {Ti}. The output from each iteration (shown in pink) serves as the input for the next iteration. (D), Average stepwise error in µm for a single embryo (e16) as a proof of concept. The black line represents the average stepwise error for 58 cells tracked over 204 minutes (35 frames), while the accumulated error is shown in pink. (E), Bidirectional approach started non-rigid registration at t(N/2), N= number of frames. (E’), Quantification of the accumulated error when the initial registration is fixed at t(N\2). (F), The comparison of average error and computation time trends of images rescaled at 10, 15, 20 and 25% of 3 embryos (e01, e02, e05). The analysis refers to the stepwise registration between only two consecutive frames. Values m1 and m2 represent the slope of the linear regressions of the two trends. (F’), The graph shows the percentage of points for which error between predicted and actual positions is greater than the mean diameter of a cell (d = 20 μm) for different rescaling rates (colour code). The dotted line represents the threshold we set to choose the resolution that best guarantees a balance between registration error and computation time. The threshold is set at 180 min of registration time. (G), Cardiomyocytes diameter is determined by calculating the maximum Feret diameter from the 3D segmentation of 26 cells. (H), The images provide a close-up view of cells under two different conditions: one where cells had already divided at time 0 (blue spots), and another where cells had not yet divided at time 0 (green spots). The orange mask outlines the segmented cell tissue at time 0 and after applying the T-transformation (t_-2h_; t_+2h_). Dotted lines illustrate the actual elongation of the tissue at t_-2h_ and t_+2h_, while the straight line within the tissue indicates the direction of maximum elongation of the transformed segmentations. (H’), Cosine similarity was calculated between the actual tissue elongation and the elongation predicted by the T-transformation for 8 cells within the -2h to +2h interval. (I), 3D image of the HT of an Nkx2.5eGFP; Rosa26Rtdtomato^+/-^ embryo(e02). The image includes a single ventral plane in a pink frame, and a single lateral plane with its segmentation section in a magenta frame. The myocardium (red), splanchnic mesoderm (green), and endoderm (blue) are visible in the segmentation section. The yellow arrow indicates the incomplete part of the right IFT. The blue arrow indicated the signal degradation in OFT. The segmentation was transformed into a volumetric mesh, defined as faces (yellow area), nodes (blue dots) and edges (orange sides). Scale bar: 100 µm. (L), Sequential interpolation between the mesh nodes with the deformation set returns a continuous tracking of the HT tissue (illustration related to e02). The points are random taken on the surface; they move in space following the tissue morphogenesis. The colour map is related to the position of the spots every hour. Yellow represents the position of the nodes at the initial time, orange after 1h, pink after 2h and purple after 3h. The grey line keeps track of the node trajectory. (M), Embryo during gastrulation. The tracking follows 25 cells for approximately 170 min. The arrows in green indicate the direction of cell motion, while the blue line tracks the entire path. (b), On the top, evaluation of the continuous error in 25 cells tracked during gastrulation at each 20 min interval. The dashed line indicates the slope (m_G_ = 3.84) of the line that fits the median error values at each interval. Scale bar: 15 µm; video time resolution: 10 min; duration of the video: 170 min. (M’), On the bottom, error related to cell tracking during HT morphogenesis at each 20 min interval. The results are relative to tracking 30 cells during 216 min. The line fitting the median values of the intervals has a slope equal to m_HT_ = 0.84. (M’’), Evaluation of the tracking error normalised for the respective cell displacements. In blue, mean values and standard deviation are shown for mesodermal cell tracking during gastrulation at 20 min intervals. In grey, values related to HT cells.

### Validating the Motion Estimation

To determine the accuracy of the algorithm in estimating myocardial motion, we used the position of randomly labelled cells in the live images as ground truth. These cells were automatically tracked and then manually corrected using Imaris software version 9.5, resulting in a total of 9584 points/cell centroids derived from 9 embryos (see Table S2). This included myocardial, splanchnic, and endodermal cells without distinction, labelled with either membrane-GFP or membrane-Tomato (Figure 2B, see also Table S2). However, only cells that did not divide during the observation period were considered for the validation process.

We created two test sets: the first set was used to estimate the intrinsic error of the registration algorithm when applied frame-by-frame (stepwise error); the second set was used to evaluate the error when the algorithm was implemented sequentially for more than two consecutive frames (accumulated error).

For the stepwise error, the test set was built applying the *morphSurf3D* function (Vedula et al., 2018), between pairs of consecutive frames. The function determined the consecutive position (x’, y’, z’) for each cell position (x, y, z) in the ground truth dataset (shown in blue/black in Figure 2C). For the accumulated error, the test set was generated by sequentially interpolating the initial cell positions (x, y, z) with the set of transformation fields {Ti}. The output of the first iteration was used as the input for the next iteration (shown in blue/pink in Figure 2C’). We measured the accuracy of the method by calculating the Eulerian distance between the position of the tracked cells (ground-truth) and the positions obtained from the registration process (test-set).

The results shown in Figure 2D demonstrate that the registration algorithm maintained a consistent stepwise error of approximately 2.5 μm throughout the video, well below the average cell diameter (∼20 μm, see also Figure 2G). However, as the registration progressed, cumulative effects and interpolation approximations ({Ti}) caused the error to increase over time. This trend is illustrated by the pink line in Figure 2D, where the mean error for a 200-minute sequence reached approximately 10 μm. To mitigate this issue, we introduced a bidirectional registration, using the midpoint of the video (t N/2) as the starting point. This divided the process into two independent registration directions (Figure 2E). As shown by the orange line in Figure 2E’, this approach successfully reduced the mean error by half. Implementing the bidirectional registration, with the starting point fixed at N/2, ensured that the maximum cumulative error remained below one cell diameter across all videos.

### Optimization Strategy

To reduce the computation time, we tested the registration process on lower-resolution images and evaluated its effect on calculation time and registration accuracy. We reduced the resolution of the images by 10%, 15%, 20%, and 25% using cubic interpolation (*imresize3* in MATLAB) and evaluated the trend of registration accuracy to determine the minimum resolution needed for acceptable results (Figure S1, STAR Methods - Optimization Strategy).

Our analysis revealed that image resolution significantly affected computation time but had a minimal impact on point registration accuracy. Figure 2F illustrates this relationship: the black line shows how computation time increased with higher resolutions (10% to 25%), while the red line indicates that tracking error remained largely unaffected by resolution changes. Based on these findings, higher resolutions were excluded from testing due to diminishing returns on accuracy.

To identify the optimal resolution, we evaluated the percentage of landmarks with errors exceeding the average cardiomyocyte diameter (20 μm) (Figure 2F’). Downscaling images to 25% resolution ensured a misregistration rate below 1% for time-lapse datasets lasting approximately 6 hours. For shorter videos (<4 hours), error for all points remained below the 20 μm threshold. Given that the average duration of 3D+t images in our study was approximately 4 hours (see Table S1, Figure S2C), this resolution was chosen.

### Continuous Description of HT Morphogenesis

The registration approach applied above captured the entire image motion, including background and other tissues. To focus specifically on the deformation of the myocardium, we segmented the tissue of interest slice by slice using three orthogonal planes in ITK-SNAP software (version 3.8.0) (Yushkevich et al., 2006). The image at time t(N/2), which is the initial registration time, was manually segmented (see STAR Methods-Image Segmentation). This resulted in a binary z stack of myocardial tissue (red mask in Figure 2I). From the binary mask, we generated a mesh using *vol2mesh* function in Iso2Mesh toolbox (Fang & Boas, 2009)(see STAR Methods-Mesh Generation). This mesh represents the surface of the myocardium, discretized into a number of points. The density of points varies for each specimen. The mesh was further refined by applying a Laplacian filter (*smoothsurf* function in Iso2mesh toolbox), to smooth its roughness (Cignoni et al., 2008). The resulting myocardial shape was called “Live-Shape”. We used the *morphSurf3D* function sequentially to move any surface mesh node at t(N/2) to its next or previous position based on the transformation set ({Ti}) creating a continuous motion profile of the myocardium surface. By iterating the process, the mesh nodes were moved in space according to the motion registered by the live image, resulting in a continuous tracking of the HT surface as shown in Figure 2L (see Video S1).

### Effects of Cell Division on Live-Shape deformation

Validation of the sequential registration process demonstrated our ability to create motion profiles with cell-scale accuracy (Figure S1D). However, during the validation process, we had to exclude all cells that underwent mitosis during video acquisition from the ground-truth. This was because the registration process assumed spatiotemporal continuity, which did not account for cell division.

To further understand the continuous behaviour of myocardial tissue in the presence of cell division, we examined the local tissue deformation and how this correlated with mesh deformation.

We then measured the correlation between the direction of cell division and the direction of deformation in the surrounding tissue. We manually segmented 8 cells before and after mitosis in a 2h registration interval. The segmented cells at t_-2h_, t_0h_ and t_+2h_ were transformed into volumetric meshes. To quantify whether the direction of elongation of the mesh captured the anisotropy generated by the final disposition of the mitotic cells, we applied the set of transformation fields to the cell nodes at t_0h_. Then, we made a line fitting of the deformed cell nodes to assess the main direction of deformation (Figure 2H, green and blue straight lines at t_-2h_ and t_+2h_ respectively). We compared this direction with the actual direction of cell division, measuring the cosine similarity between the two directions. The results demonstrated that the tissue exhibited elongation in the direction of cell division with an average cosine similarity value of 0.87 (SD=0.11) for cells at +2h and 0.87 (SD=0.10) for cells at -2h (Figure 2H’). This indicates that the image-based algorithm captures the change in geometry and directionality resulting from cell division, which further supports the accuracy of the proposed method.

### Limitations of the approach

The continuous description employed did not allow the exchange of mesh node positions. Consequently, the observed registration accuracy at cellular level suggests that during heart morphogenesis, cells largely maintain their relative positions within the timescales used in our analyses. To test this idea, we studied the performance of the procedure in a tissue known to show high cell mixing: the nascent mesoderm during gastrulation. We applied the registration approach to 25 mesenchymal cells during gastrulation (see STAR Methods - Embryo culture and live imaging of gastrulating mouse embryos). At this stage of embryogenesis, cells in the mesoderm exhibit strong mixing behaviour (Ichikawa et al., 2013), resulting in a more chaotic pattern of movement (Figure 2M). We compared tracking error in 170 min with the accuracy of the algorithm when applied to early heart tissue morphogenesis (Figure 2M’). This analysis utilized the live image data of embryo e02. The results demonstrate that the algorithm achieved an accuracy significantly below 20 µm for the initial 20 minutes of recording when applied to gastrulating cells. However, after this initial period, the accumulated error increased approximately five times faster for mesenchymal cells during gastrulation than for cardiomyocytes during heart morphogenesis (m_G_/m_HT_, where m_G_ = 3.84 is for gastrulation and m_HT_ = 0.84 is for HT).

This comparison highlights the limitations of the algorithm when applied to uncoordinated cell movements and emphasizes the highly coordinated dynamics and low mixing behaviour of cardiomyocytes. Mesodermal cells, with their mixing behaviour, exhibited less displacement within the tested time window compared to cardiac cells. Myocardial cells displayed higher straightness and moved more cohesively. The estimated tracking error, normalized by cell displacement, demonstrated a significant difference in registration pipeline performance between the two cases (Figure 2M’’). This illustrates that the algorithm was more effective in tracking coordinated, low-mixing movements as opposed to uncoordinated, and chaotic movements.

### Integrating Multiple Live Images into a Consensus Temporal Reference

To compare different specimens and capture the intrinsic variability, it is essential to fix a spatio-temporal reference (see Video S2). To address this challenge, firstly, we have developed a staging system that aligns live images of different specimens over time based on a morphometric feature. Then, we have designed a strategy to overcome the shape variability between equally staged hearts by projecting all specimens onto a unique spatial reference system. We have designed a systematic approach for synchronizing live images in a common temporal framework. To achieve this, we have employed the Atlas proposed Esteban et al., as a temporal reference system (Esteban et al., 2022). The Atlas offers a comprehensive and detailed description of the morphology of the heart tube at tissue level during the development process (E7.75-E8.5), from the cardiac crescent to heart tube looping. Esteban et al. discretized this developmental window into 10 groups using a morphometric parameter, d_1_/d_2_, to cluster the shape of 50 specimens (Figure 3A). This parameter is the ratio between the length of the boundary between the myocardium and the juxta-cardiac field (d_1_) and the length between the myocardium and the splanchnic mesoderm (d_2_) (as depicted by the green line in Figure 3A).

**Figure 3:**
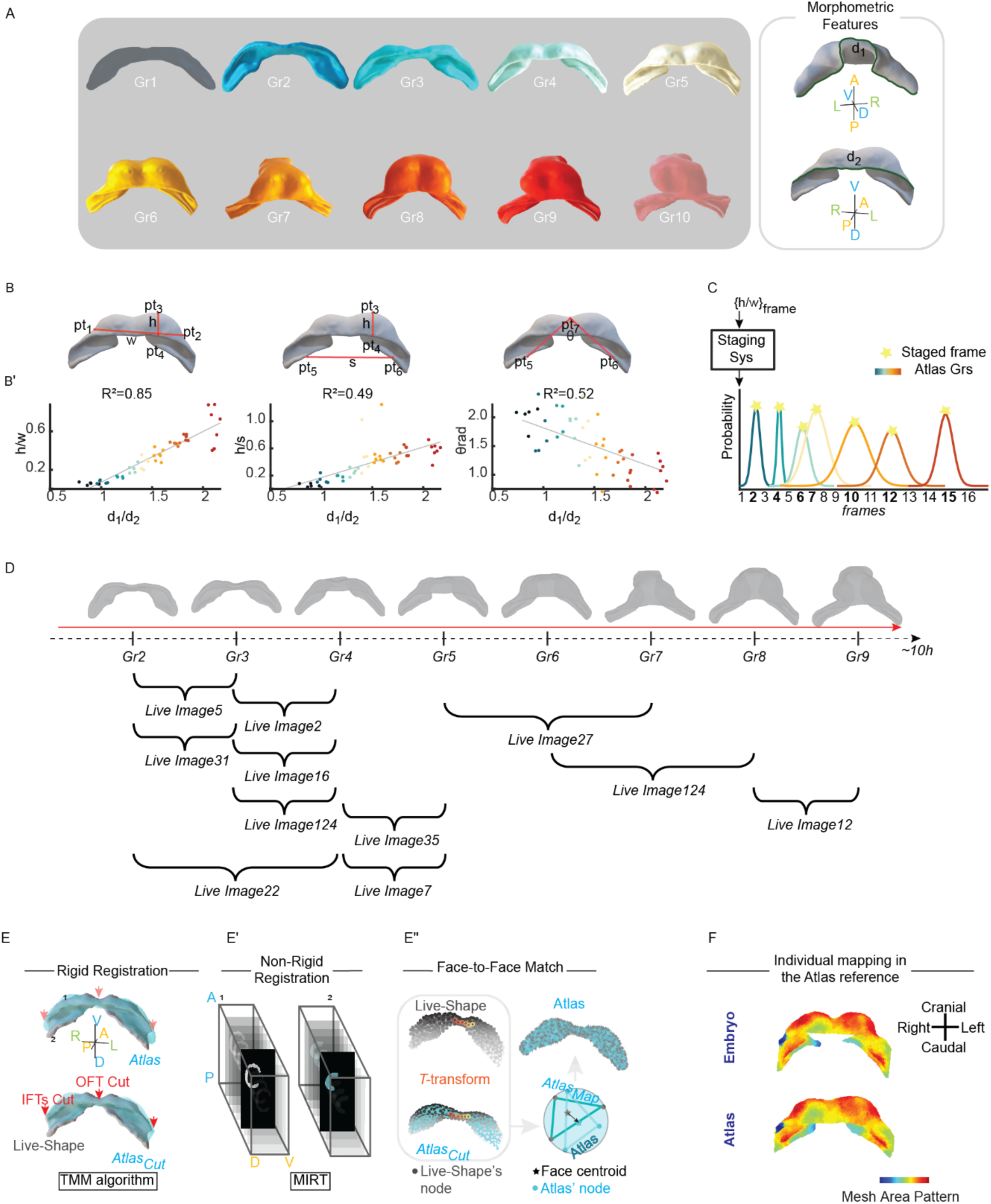
Integrating multiple live images into a consensus spatiotemporal reference by aligning each continuous motion profile with a published 3D+t Atlas. (A), The static Atlas is described by 10 representative heart shapes. On the right, the morphometric parameter d1/d2 is shown. (B), Proposed features are defined as the Eulerian distances h, w, s, and θ. (B’), From left to right, a linear regression profile is shown for the ratios h/w, h/s, and θ versus d1/d2. Each scatter plot displays all Atlas specimens belonging to groups 1 to 9 (48 out of the total 50 specimens). The different groups are represented by a colour code. For each scatter plot, the coefficient of determination (R²) is reported. The h/w ratio has the highest correlation value with d1/d2, with R² = 0.85, while the linear regression models for h/s and θ returned R² values of 0.49 and 0.52, respectively. (C), Staging System: A Gaussian Mixture Model (GMMs) associates to the Atlas groups (indicated by colour code) the live image frame with highest probability (denoted by stars). (D), Results of the staging system. (E), TMM rigid registration rotates, translates, and resizes the Atlas shape (blue surface) until it overlaps with the Live-Shape (grey surface). Parts of the Atlas shape corresponding to the missing IFTs and OFT in the Live-Shape (indicated by red arrows) are removed, resulting in the Atlas_cut_. (E’), Image non-registration (MIRT) is used to transform the Live-Shape mask to the Atlas_cut_ mask. (E’’), The T-transformation adjusts the position of the Live-Shape nodes (grey point-cloud) to fit the Atlas_cut_ (blue point-cloud) morphology. The morphed shape is called SurfaceMap. The coloured points indicate the same nodes in their configuration before and after the transformation. Face-to-face matching between the Atlas shape and the Live-Shape is performed between the centroids of the faces (marked by stars). (F), Validation of the spatial mapping is conducted by computing the mesh area in the Live-Shape and plotting the growth values onto the Atlas face-to-face.

The lengths, d_1_ and d_2_, were not measurable on Live-Shapes due to missing parts; specifically, the IFTs and OFT as seen in Figure 2G (indicated by the yellow and blue arrow). To overcome this challenge, we studied alternative features that would produce similar staging results than d1/d2. We introduced three morphometric features h/w, h/s, and θ (Figure 3B). These features were computed by fixing 7 specific landmarks on the Live-Shape (STAR Methods - Selecting Myocardial Landmarks). The first two features, h/w and h/s were defined as the ratios between the Eulerian distances between the landmarks, while the third feature, θ, was calculated as an angle value. We excluded Atlas group 10 from our staging system as the live images collection did not cover the heart looping stage. Therefore, we calibrated our new staging system using 48 of 50 samples from the Esteban et al. collection (E7.75-E8.25 12h). To define the three features, we manually selected the 7 landmarks (pt_1_-pt_7_) on the mesh of these specimens. To minimize the manual error, we selected the 7 landmarks three times, and we took the median landmark coordinate values.

Once the 7 landmarks on the 48 shapes had been selected, we calculated the Euclidean distances between pt_1_ and pt_2_ (w) and between pt_3_ and pt_4_ (h). Furthermore, we defined θ as the angle formed between pt_7_ and the points pt_5_ and pt_6_. These measurements showed a monotonic trend over time, which allowed to correlate their value with the developmental stage (Figure S3A,B). To normalize the data and control for variations in myocardial size between specimens, we defined the morphometric features by taking ratios between these measurements, except for θ, which needed no correction for the size of the myocardium.

Subsequently, we evaluated which of the proposed features had the strongest correlation with d_1_/ d_2_ by performing a stepwise regression (*stepwiselm* function in MATLAB) (Draper & Smith, 1998). The result showed that the proportion of h/w exhibited a high degree of correlation with the d_1_/d_2_ parameter, with a coefficient of determination (R^2^) of 0.85 and a p-value of 2.30e-20 (Figure 3B’). Given the strong correlation between h/w and d_1_/d_2_ (up 85%), we used this feature to stage the Atlas for the purpose of directly staging the live images with respect to the Atlas reference.

We then modelled the staging system as a Gaussian Mixture Model (GMMs) being the h/w values normally distributed for each Grs (STAR Methods-Modelling the Staging System; see also Table S4). Then, we collected h/w distances for each Live-Shape extracted from the 3D+t images. The landmarks pt_1_, pt_2_, pt_3_, pt_4_ were fixed only on the Live-Shape relative to the frame (N/2). The landmarks positions for the remaining time-points were determined based on the continuous motion profile. This ensured that we were able to optimize time, but also to mitigate any potential bias that may have been introduced by manually selecting the landmarks (Figure S3).

The classifier returned for each image frame the probabilities of belonging to Atlas groups. The belonging value closest to 1 determined the time-matching (Figure 3C). Since the temporal resolution of the live imaging is higher than the pseudo-temporal resolution of the Atlas, several time-lapse frames were associated with the same Atlas reference (Figure 3D). To handle this issue, we selected only the frame with the highest probability among the equally clustered frames as the representative frame (Table S6).

### Registering Individual Shape into the Atlas Reference Geometry

We considered the Atlas not only as a temporal reference, but also as a spatial reference. So once a shape was staged, that shape was projected into Atlas, which allowed a single geometrical representation for otherwise variable shapes. With this aim, we implemented a mapping pipeline that combines rigid and non-rigid registration techniques in three steps: 1. Rigid registration of Atlas shape with the Live-Shape; 2. Non-rigid registration between Live-Shape mask and the Atlas mask; 3. Face-to-face matching between the registered Live-Shape in the corresponding Atlas shape.

The first step involved the rigid registration of Atlas to the Live-Shape (Figure 3E; see also Table S5). We employed a [TMM-based algorithm](https://es.mathworks.com/matlabcentral/fileexchange/63693-robust-group-wise-registration-of-point-sets-using-multi-resolution-t-mixture-model?s_tid=prof_contriblnk) (Ravikumar et al., 2016, 2018)(see STAR Methods - TMM setting parameters), which leverages the robustness of Student’s t-distributions against outliers. This approach enhanced the overlap results, even in the presence of missing regions in the Live-Shape. Once the two heart shapes were rigidly superimposed, we imported them into MeshLab and eliminated from the Atlas shape all nodes for which there were no correspondence with the Live-Shape, resulting into the Atlas_Cut_ version (Figure 3E; STAR Methods - Removing Missed Correspondences).

The second step was aimed at identifying the necessary transformation to morph the Live-Shape into the Atlas_Cut_. This was accomplished by performing a non-rigid registration in the image domain. We implemented the MIRT algorithm between the Live-Shape mask and the Atlas_Cut_ mask (Figure 3E’; STAR Methods-Defining the Live-Shape Mask, Defining the Atlas Mask). The resulting T-transformation was applied to the Live-Shape, modifying its nodes arrangement according to the Atlas_Cut_ surface. This modified version of Live-Shape is called SurfaceMap, describing the anatomical correspondences between the Live-Shape and the Atlas_Cut_. For each Gr, we computed the SurfaceMap of staged specimens. These SurfaceMap, although describing the same Atlas surface, have a different node density. The sequence of SurfaceMap of the same specimens in sequential Atlas stages describe its individual profile motion in the Atlas reference. The last step of the spatial mapping resolved node density variability by uniquely finding correspondences between SurfaceMap and Atlas. We associated the centroid of each Atlas face to the nearest SurfaceMap centroid face (*knnsearch* function in MATLAB), resulting in a face-to-face matching between the two shapes (Figure 3E’’). Since there was no numerical way to test the accuracy of spatial mapping, we opted to assess its efficacy by examining the growth colour pattern. We utilized the *meshFaceAreas* function in the [geom3D library](https://github.com/mattools/matGeom/wiki) to calculate the area values of each Live-Shape mesh triangles. We plotted the area values on the Atlas in locations where face-to-face correspondences were identified. We observed that the colour patterning between the Live-Shape and the Atlas (Figure 3F) was consistent.

### Extracting Stepwise Tissue Deformation

In the continuous description of the HT morphogenesis, mesh deforms according to *T-* transformations; nodes move, and edges change their configuration, generating an effective deformation of each face (Figure 4A). We computed these mesh deformations by using the continuous mechanics laws between the rest state and the deformed state (Lai et al., 2009; STAR Methods - Principles of Finite Deformation Continuum Mechanics). Changes in mesh face configuration were quantified as tissue growth rate (J) and tissue anisotropy (θ).

**Figure 4:**
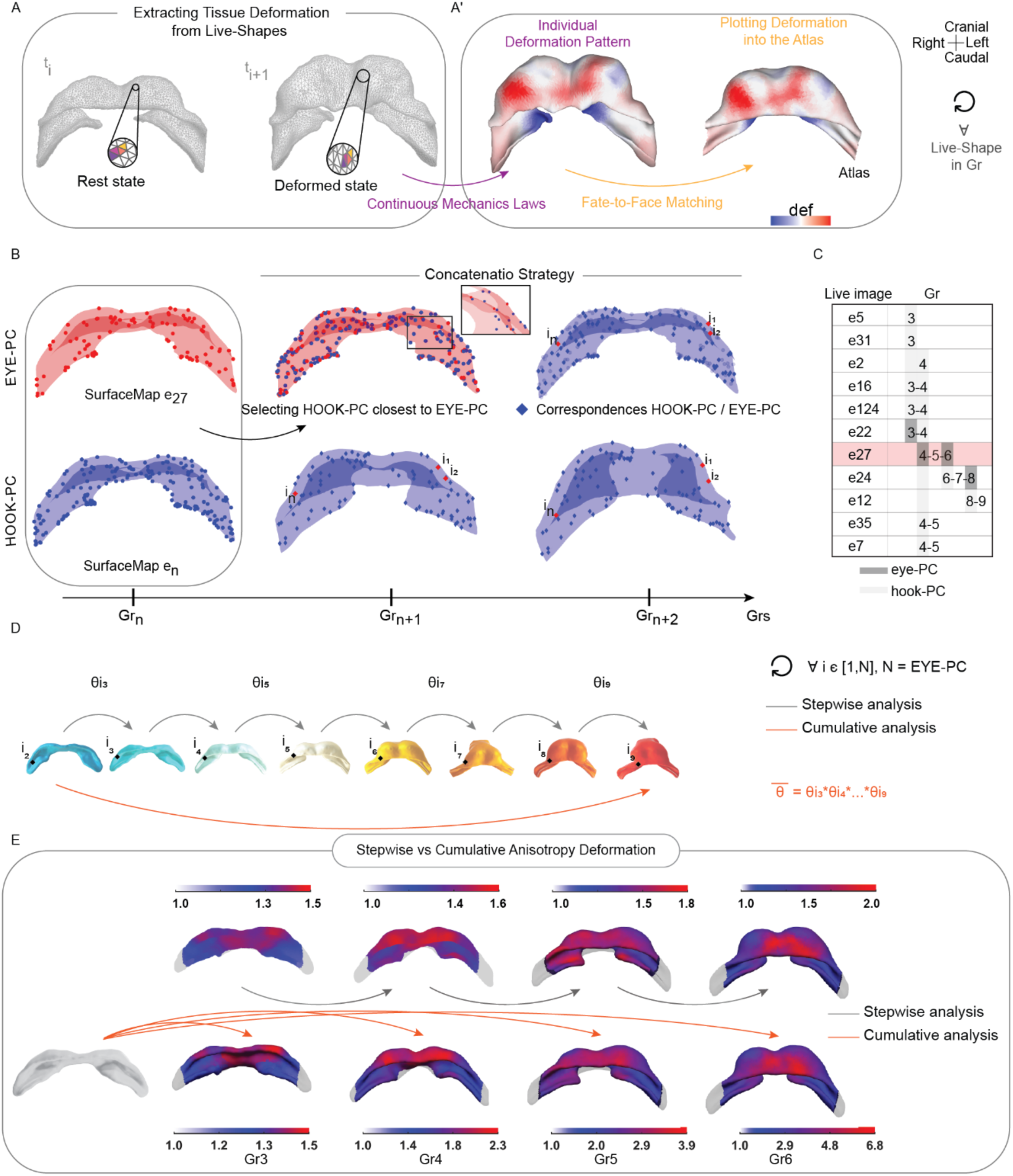
Quantifying tissue deformation during early morphogenesis. (A), The triangles of the mesh were transformed according to the T-transformation from t_i_ to t_i+1_. Colours code indicated the same triangles in their rest and deformed state (shapes refer to e27). (A’), The tissue deformation pattern into is plotted in Live-Shape(t_i+1_) and mapped face-to-face into the synchronise Atlas group. We compute the individual deformation pattern for each staged Live-Shape and for each Gr. (B), Schematic overview of the pipeline for concatenating multiple motion profiles using the EYE-PC and HOOK-PC strategies. SurfaceMap e27 at Gr_n_ represents the EYE-PC (red shape and dots). To predict the position of SurfaceMap e27 in subsequent groups (Gr_n+1_, Gr_n+2_), we used known SurfaceMap from another specimen, referred to as the Hook-PC (blue shape and spots). The closest points in the HOOK-PC to the EYE-PC at Gr_n_ were selected as corresponding points (blue rhombuses). The position of SurfaceMap e27 (i_1_, i_2_, …, i_n_) in Gr_n+1_ and Gr_n+2_ was then determined by the positions of the selected points from the Hook-PC. (C), Live images concatenation path. Embryo e27 is the reference. In dark grey the eye-PCs are highlighted, in light grey the hook-PCs. (D), The cumulative anisotropy rate (*θ̅*) of each mesh face (i) between two arbitrary Grs is given by the product of all the intermediate *θ̅*_gr_. (E), Stepwise vs Cumulative anisotropy (*θ̅*). Cumulative deformation is computed from Gr_2_. Colour bars indicate the deformation magnitude. *θ̅* = 1 isotropic deformation. *θ̅* > 1anisotropic state. Caudal view of the heart tube is shown.

We extracted tissue deformation for each live specimen with at least two frames classified in consecutive Atlas Grs (12 embryos, see Table S6). The deformation pattern was always plotted into the shape of the end frame. Gr_1_ was excluded from the deformation analysis, as our focus was on quantifying tissue changes directly related to morphogenesis events, whereas from Gr_1_ to Gr_2_ the main event is differentiation, not morphogenesis.

To compare the deformations between individual specimens at the same stage, we plotted the deformation onto the Atlas shape using the face-to-face matching (Figure 4A’). To investigate the spatial consistency of deformation patterns across specimens, we performed the mean deformation (µ_def_) and standard deviation (σ_def_) within each group.

### Calculating Cumulative Deformation

The Atlas does not offer any kinetic data since the mesh nodes lack correlation between Groups. Therefore, to calculate the cumulative deformation of the tissue starting from Gr_2_ and ending in Gr_9_, we deduced the Atlas surface motion. To achieve this, we developed a strategy that concatenates individual motion profiles, allowing us to track tissue deformation over time. The concatenation was carried out using the SurfaceMaps of the staged specimens (see STAR Methods - Concatenating Individual HT Motion Profiles to Compute Cumulative Deformation). The SurfaceMaps represented the motion profile of each specimen within the Atlas. To merge multiple SurfaceMaps, we addressed the varying point densities by selecting one SurfaceMap density distribution as reference (e27). This choice was made because e27 live images encompassed a central portion of the developmental timeline, specifically Gr_4_, Gr_5_, and Gr_6_. We estimated how the e27 SurfaceMap deformed before Gr_4_ and after Gr_6_ by using SurfaceMap_s_ from specimens at earlier or later stages (Figure 4B). We found correspondences between the e27 SurfaceMap Point-Cloud (eye-PC) and those of other specimens (hook-PC) (Figure 4C). This enabled us to generate a continuous motion profile, allowing the computation of the cumulative deformation of the developing tissue.

For each group, we computed the mean growth rate 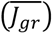 and the mean anisotropy value 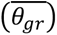 based on the i-correspondences between eye-PC and hook-PC. Then to determine the cumulative tissue growth (*J*) from Gr_2_, we multiplied the mean growth rate values 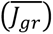 at each *i*, across all groups (as illustrated in Figure 4D):

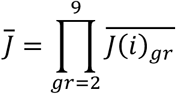

where 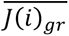 is the stepwise mean growth rate for each Gr. Similarly, we analysed the cumulative anisotropic deformation *θ̅*, by calculating:

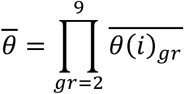

where 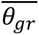 is the stepwise mean anisotropy computed. In Figure 4E, the first row illustrates the cumulative growth of the myocardium. The red areas (*J* > 1) represent tissue expansion, primarily concentrated on the lateral side of the rounded chamber. The white areas (*J* = 1) indicate regions where the tissue size remains unchanged, such as the medial region of the chamber. The blue areas (*J* < 1) signify tissue compression. In the second row, the image shows the anisotropic behaviour of the deformation. Areas of strong anisotropy are shown in red (*θ̅* >>1). White areas represent isotropic deformation (*θ̅* =1).

### In-silico Fate Map

We employed the concatenation strategy to develop an *in-silico* fate map of cardiomyocytes during early heart morphogenesis. By tracking a single SurfaceMap from Gr_2_ to Gr_9_, we created a dynamic Atlas (see STAR Methods - Concatenating Individual HT Motion Profiles to Compute in Silico Fate Map). This dynamic Atlas provides crucial insights into potential cell rearrangements within the myocardium. Assuming each point in the cloud represents a myocardial pseudo-cell, we were able to simulate tracking of each pseudo-cell from Gr_2_ to Gr_9_. To achieve this pseudo-cell tracking, we chose not to combine the kinetics of multiple embryos per group but, instead, concatenate single motion profiles for each Gr. This approach allowed us to limit the artifacts related to missing parts of IFT and OFT. Since these missing parts are not systematic, we had to find the common part of the heart included in all considered specimens and cut out the uncommon parts. To create the fate map, we established a concatenation path illustrated in Figure 5A, which involved using as many live images as possible, therefore we used e31, e16, e35, e27, e24, e12 (Figure S4). The resulting dynamic Atlas maintained the original morphology of the static version (Esteban et al., 2022), but the positions of the mesh nodes followed the kinetic derived from the concatenation of the motion profiles of myocardial cells in the live embryos (Figure 5B, Video S3).

**Figure 5:**
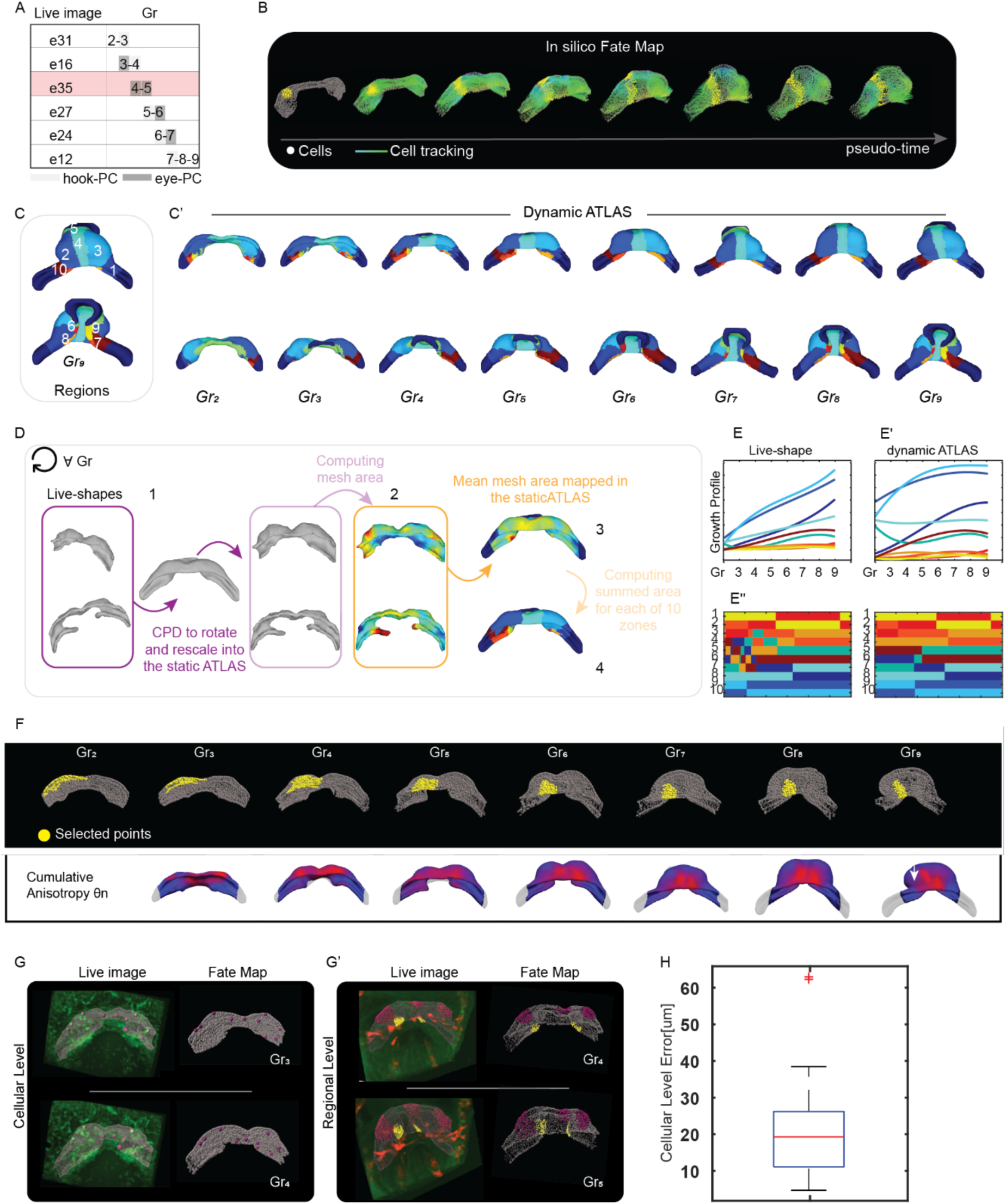
Creating an *in-silico* fate map to analyse heart tube morphogenesis at both cellular and regional levels. (A), Concatenation path of the individual live image. We concatenated through the hook-PC and eye-PC strategy the videos related to embryos e31, e16, e35, e27, e24 and e12. The reference embryo is e35. (B), *In-silico* fate map tracking early cardiac morphogenesis as a continuum. The tool in Imaris allows to select points or regions in the myocardium and follow their motion over time from Gr_2_ to Gr_9_ in Atlas. In grey the point-cloud, in yellow the selected points track the regional changes at different stages. The lines, on the other hand, draw the displacement of the point-cloud. (C), The selected 10 arbitrary zones of the myocardium on the Gr_9_ shape. (C’), The displacement of the 10 zones in the different groups of the dynamic Atlas. The two rows above represent the anterior part of the heart, the two rows below the posterior part. (D), Strategy to identify the same 10 anatomical zones in the live embryo. For each group, Live-Shapes were rescaled with a rigid registration algorithm to the Corresponding Atlas shape. Then the area of each mesh was calculated (colour map) for each rescaled Live-Shapes. Finally, the area values were averaged within the Atlas reference and summed for each of the 10 areas. (E-E’), The Live-Shape and the dynamic Atlas growth profiles of the 10 zones, interpolated for each Grs with a 3-degree B-Spline. (E’’) The validation strategy. (F) Top: A selected region in the dynamic Atlas, illustrating how the region deformed anisotropically over time. Bottom: Cumulative anisotropy from G_r3_ to Gr_9_. (G), At the cellular level (left), we tracked cell positions in e02 at Gr_3_ and Gr_4_ using Imaris (pink spots). On the right, we marked the initial positions of these cells on the fate map at Gr_3_ and then evaluated their predicted final positions at Gr_4_ (pink spots). (G’), At the regional level, we tracked different groups of points on the myocardium mesh of e27 from Gr_4_ to Gr_5_ (pink and yellow points). On the right, we identified the same groups of points on the fate map at Gr_4_ and assessed where these points ended up in Gr_5_. (H), Cellular-level error across 40 tracked cells from e01, e02, e05, and e16, compared to the corresponding predicted positions from the dynamic atlas. On average, the error is 20 µm, roughly equivalent to a cell diameter. Only 2 cells show an error greater than twice the cell diameter. We consider 95% of the cells to be well-predicted.

### Validating the Fate Map Deformation

To evaluate how well the fate map predicts myocardial deformation, we validated that the dynamic Atlas motion profile accurately reproduced myocardial growth. We compared the average growth patterns of 10 selected zones in live specimens with those in the dynamic Atlas. These zones were identified in the dynamic Atlas at Gr_9_ (as shown in Figure 5C) and tracked across other Grs (Figure 5C’).

To determine the dynamic Atlas growth profile, we calculated and smoothed the mesh areas for each zones using the *meshFaceAreas* function in the [MatGeom](https://github.com/mattools/matGeom/wiki) toolbox. For each zone, we summed the area values defining the Atlas growth profile, shown in Figure 5E’ as three-degree B-splines. To determine the Live-Shapes growth profile we identified the same 10 zones as illustrated in Figure 5D. We first rescaled each specimen to the reference static Atlas shape using the Coherent Point Drift (CPD) algorithm (Myronenko et al., 2007)(step 1 in Figure 5D). We then computed and smoothed the mesh areas for these rescaled Live-Shapes (step 2 in Figure 5D). We mapped and averaged the individual mesh areas onto the static Atlas (step 3 in Figure 5D). As last step, we found the 10 zones as the correspondences between the static Atlas and dynamic Atlas (step 4 in Figure 5D). The Live-Shape growth profile is shown in Figure 5E’.

We compared the growth profiles by checking the consistency at each x-point (Figure 5E’’). We verified this consistency by correlating the sorting growth values of each zone between the two profiles across the groups (Figure 5E’). This method allowed us to examine myocardial growth profiles across different heart zones, independent of growth magnitude, while accounting for embryonic variability. Our analysis found that the growth profile of the dynamic Atlas matched that of the Live-Shapes with 81% accuracy (see also Figure S5). The error rates for zones 1, 10, and 5, corresponding to the left and right IFT and OFT, respectively, were relatively high due to manual cutting and misalignment issues during spatial mapping. Removing these zones improved the motion evaluation accuracy to 92%. At the regional level, we also evaluated the tissue deformation. Clearly, the ventral part of the growing ventricle suffered deformation in the craniocaudal direction (Figure 4E). Similarly, the la fate map deformed anisotropically in the medial area, as shown in Figure 5F.

### Validating the Fate Map Motion

We then performed virtual labelling of single cells or regions within the dynamic Atlas and compared the fate of the labelled regions with tracking of the same positions directly in the live image, which were considered as ground truth (Figure 5G-G’). The accuracy of the Dynamic Atlas was quantified by tracking the positions of 40 cells from their initial to final position in live images. Subsequently, we compared the tracked final position with the predicted final positions provided by the fate map. Given the intrinsic variability of the shapes, we employed a manual approach to validate the accuracy of the predicted positions. We defined a small surrounding area (∼ 2 cells diameters) on the fate map within which we expected to find the point at its final position. If the predicted position fell within this defined area, we considered it to be correct. We found that the location of 95% of the selected points/cells was correctly predicted according to the validation criteria (Table S7).

## DISCUSSION

We designed and developed an image-based pipeline to extract and compare tissue motion and deformations during early morphogenesis and applied it here to mouse heart development. Our method is proposed as a computational solution to address biological challenges and overcome technological limitations in mammalian organogenesis. This approach enables the non-invasive extraction of myocardial motion and deformation, successfully managing the intrinsic variability of the phenomenon. To the best of our knowledge, the only method delivered for the extraction of the cardiac tissue deformation map at cellular resolution during morphogenesis was designed by Kawahira et al., 2020 for the C-looping in chicken heart. In that case, the authors quantified myocardial torsion deformation from the tracking of several hundred cardiomyocytes.

Such a strategy would not be feasible for the drastic deformations that many tissues undergo during organogenesis, as it is the case for heart tube formation from the cardiac crescent. In fact, early cardiac morphogenesis involves complex shape changes, including changes in topology, and a detailed reconstruction of deformation maps would require over thousands of tracked cells. But it is difficult to automatically track cells in such a dense population. This difficulty is mainly related to the limited spatial and temporal resolution of the mouse heart images and the variation in fluorescence intensity caused by photobleaching. Our strategy takes advantage of the intensity of the image and, as demonstrated, does not require the high resolution required for cellular tracking. The validation process showed that the configured registration method succeeds in tracking cell motion profile with cell-scale accuracy (Figure 2F’, Figure S1). Despite the continuity assumption, the method was successful in tracking cell division at the tissue level, indicating finally that the implemented data-driven approach was efficient in the characterization of early heart tube morphogenesis. Interestingly, the method revealed the coordinated nature of cardiomyocyte motion, denoting moderate cell mixing during the shaping process (Figure 2M-M’’).

To compare different specimens, we needed to deal with the intrinsic variability of HT morphogenesis. To combine and compare multiple kinetic data, it was necessary to define a common reference system. We chose the Atlas shapes as our spatio-temporal template (Esteban et al., 2022). The Atlas not only narrows down the number of developmental stages to be considered, but also solves the problem of incomplete shapes, which poses an added degree of complexity.

We have introduced a standalone morphometric staging system that enables the synchronization of live images along the Atlas temporal axis, utilizing a simple geometric feature. This feature, defined as the ratio of two Eulerian distances within the primitive chamber of each Live-Shape, offers a significant advantage: it can be applied directly to 2D images, bypassing the need for complex 3D reconstructions. In contrast, the staging system proposed by Esteban et al., 2022 relies on a geometric parameter (d1/d2) derived from geodesic distances, which necessitates a full 3D reconstruction of the myocardium and a complete capture of the heart. This requirement poses a challenge for live imaging, where depth resolution is often limited, and anatomical regions can be lost due to partial movement of the embryo out of the field of view. To validate our staging system’s effectiveness in aligning live images with the Atlas timeline, we performed a statistical analysis to assess the correlation between our geometric feature (w/h) and the d1/d2 parameter. The results demonstrated a strong correlation (Figure 3B’), indicating that the w/h feature is a robust alternative for mouse heart staging, consistent with the temporal discretization provided by Atlas.

To address variability in myocardial shapes of the same stage, we implemented spatial mapping using registration techniques to adapt the Live-Shape anatomically to the Atlas. Non-rigid image registration (Myronenko et al., 2007) proved more efficient and robust than point cloud-based registration, which struggles with the complex and variable myocardial shapes found across specimens. Point cloud-based methods are limited by their reliance on a small number of surface nodes, often failing to find accurate matches, especially on the inner and outer sides of thin heart tissue. To overcome these limitations, we converted point-clouds into binary image masks, enabling the matching of a significantly greater number of anatomical points (each voxel) and ensuring accurate correspondences on the correct side of the tissue. Although there is no analytical method to directly measure spatial mapping accuracy, the mesh area pattern from the Live-Shape was precisely mapped onto the Atlas shape (Figure 3F). In fact, the colour pattern consistently matched the Atlas shape, thereby establishing a valid correspondence between the Live-Shape and the Atlas.

Deformation patterns were derived from the motion profile of each specimen in the live collection. Comparing these patterns across specimens required addressing the inherent variability of heart tube morphogenesis. To identify differences and extract common features, a standardized reference system was needed. We selected Atlas shapes as the spatio-temporal reference, focusing specifically on Gr_2_ through Gr_9_ as our reference stages. We computed the cumulative deformation that occurred from one stage to the next, capturing the progressive changes in heart tissue from Gr_2_ to the subsequent stages. The innovation of the proposed cumulative method mainly concerns the possibility of recreating a complete deformation profile starting from the heart primordium by linking multiple live images. Obtaining live images that span throughout entire early heart development is challenging and requires high-performance microscopy. The implemented cumulative strategy allowed for the computation of deformations between any selected stages, providing flexibility. Here, we have chosen to describe the transformation that the Gr_2_ shape undergoes until heart tube formation.

By integrating motion profiles from multiple specimens, we have created the first *in-silico* fate map for early heart tube morphogenesis, establishing a dynamic Atlas. This digital motion profile accurately reproduces the natural growth of the heart tube, which we explored here in 10 distinct arbitrary regions, thereby demonstrating the reliability of the fate map. Furthermore, the fate map reproduces the anisotropic deformation of the tissue, especially in the central region, where the craniocaudal cumulative deformation analysis suggests the greatest anisotropy. Locally, we validated the fate map’s predictive accuracy by tracking 40 cell positions on live images, showing that the map effectively models cardiomyocyte positions across stages.

Based on these results, we propose the fate map as an interesting tool for selecting points or theoretical myocardial cells (pseudocells) and tracking their movements from Gr_2_ to G_9_. This tool not only provides a novel approach for exploring cardiac morphogenesis at the cellular and regional level but also presents an innovative way to conduct motion analysis. This innovative tool sheds light on regionalized cell coordination, generating new questions about the biological and genetic factors that regulate the cardiac development process. Beyond cardiac development, the methods and concepts described in this work can be generally applied to the study of embryogenesis and organogenesis.

## Materials and Methods

### Data and code availability

The authors declare that all data supporting the findings of this study are either included in the article and its supplementary information files or can be obtained from the corresponding author upon reasonable request. Additionally, all raw and processed data generated during this study have been archived on a dedicated Mendeley Data server and can be accessed at the following addresses:

- Mendeley Data: https://data.mendeley.com/preview/54gbvnsgnp?a=6dc681ed-ea8c-4e15-bc00-dcc62b3a6178
- DOIs are listed in the key resources table.
- Code is available in https://github.com/MorRaiola/QuantitativeAnalisysOfHTMorpho-genesis.git

### Experimental workflow

#### Mouse Strains

Animals were handled in accordance with CNIC Ethics Committee, Spanish laws and the EU Directive 2010/63/EU for the use of animals in research. All mouse experiments were approved by the CNIC and Universidad Autónoma de Madrid Committees for “Ética y Bienestar Animal” and the area of “Protección Animal” of the Community of Madrid with reference PROEX 220/15. For this study, mice were maintained on mixed C57Bl/6 or CD1 background. We used the following mouse lines (Table S1), which were genotyped by PCR following the original study protocols. Male and female mice of more than 8 weeks of age were used for mating.

#### Embryo Culture and Live Imaging of Gastrulating Mouse Embryos

Live imaging procedures followed the protocol outlined in Sendra et al., 2022. In brief, mouse embryos were carefully collected and dissected within a dissection medium comprised of DMEM supplemented with 10% fetal bovine serum, 25 mM HEPES-NaOH (pH 7.2), and penicillin-streptomycin (50 µg/ml each). For embryos spanning E6.5 to E7.5, culture conditions were established using a mix of 50% Janvier Labs Rat Serum Sprague Dawley RjHan SD (male only) and 50% DMEM FluoroBrite (Thermo Fisher Scientific, A1896701) with incubation at 37°C and a 7% CO2 concentration. Imaging was conducted on a Zeiss LSM780 platform, featuring a 20× objective lens (NA=1) and a MaiTai laser set at 980 nm for two-channel two-photon imaging. Fluorescence was detected with Non Descanned Detectors equipped with the filters cyan-yellow (BP450-500/BP520-560), green-red (BP500-520/BP570-610) and yellow-red (BP520-560/BP645-710). Zen software (Zeiss) facilitated data acquisition with an output power of 250 mW, pixel dwell time of 14.8 s, line averaging of two, and an image dimension of 1024×1024 pixels.

### Computational workflow

In this section, we provide a detailed explanation of the steps within the framework. We outline the computational strategy employed to extract, measure, and compare deformation patterns from 3D time-lapses, by integrating image processing, computer vision, statistical analysis, and physics concepts.

#### Image Preprocessing

The raw images were captured using a two-photon microscope (Zeiss LSM780) (Ivanovitch et al., 2017). Some images were significantly affected by shift. To address this, we applied block-matching registration to correct for both translational and rotational misalignments across all planes, using the [registration tool](https://github.com/GuignardLab/registration-tools) developed by McDole et al. (McDole et al., 2018). Most images included two reporter alleles, corresponding to two channels (ch1 and ch2). To integrate the information from both channels, we combined ch1 and ch2 using Fiji (Schindelin et al., 2012) to produce a single-channel image, I(x, y, z, t). This combined image was then imported into MATLAB via the JAVA MIJ package MIJ (*ij.IJ.openImage* function, *ImagePlus2array* function) (Giovannucci et al., 2019; Sage et al., 2012), and cropped using the *imcrop3* function to eliminate non-cardiac tissue and reduce the image size. In the raw data, voxels had varying dimensions along the x, y, and z axes (anisotropic voxel [aμm; aμm; bμm]). To ensure comparability across all dimensions, we resampled the images using nearest-neighbour interpolation, converting the anisotropic voxels into isotropic voxels with equal dimensions along all three axes [aμm; aμm; aμm]. This resampling was performed with the *imresize3* function. To reduce noise, we applied a 3D Gaussian filter (*imgaussfilt3*) with a sigma value of 0.5 as needed. Next, we scaled the intensity range of each 3D image to [0-1] using the *mat2gray* function to mitigate the effects of photobleaching. Finally, we rotated the hyperstacks by 90° around the y-axis in the transverse plane using the reslice function in Fiji. The processed hyperstacks were then saved in the Tagged Image File Format (TIFF).

#### MIRT algorithm

In MIRT (Medical Image Registration Toolbox) non-rigid registration algorithm (Myronenko et al., 2007; Myronenko & Song, 2010; Song et al., 2006), the alignment between two images, I and J, is accomplished by minimizing a target function, E_target_. This target function is composed of three key components: the similarity measure (E_sim_), the transformation model T(x), and the optimization method:

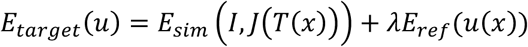

As similarity function E_sim_, we have decided for the Sum of Squared Differences (SSD).

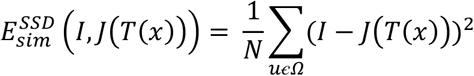

where *N* is the total number of voxels. *E_sim_^SSD^* quantifies the intensity difference for each voxel position in the 3D images domain Ω. SSD is one of the simplest intensity-based similarity measures. Nevertheless, it guaranteed good results when applied to live heart images. This was because consecutive images exhibited a high degree of similarity, owing to the temporal resolution being sufficiently high.

As transformation function T(x), the algorithm uses Free-Form Deformation (FFD) transformation (Sederberg & Parry, 1986), which generates smooth deformations. This transformation model parameterized deformations on a per-pixel basis, i.e. it deforms a 3D image (Ω = (x,y,z)| 0 ≤ x_n_ ≤ N, 0 ≤ y_m_ ≤ M, 0 ≤ z_k_ ≤ K) by manipulating a regular grid of control points (blue grid showed in Estimating Individual Live Image Motion in Figure 1) that are distributed across the image at an arbitrary mesh resolution (nx x ny x nz). The spacing of the regular grid points governs the transformation scale. At each iteration, the control points update their position (orange grid showed in Estimating Individual Live Image Motion in Figure 1), and a dense deformation is computed by using a B-spline transformation (De Boor & De Boor, 1978).

We fix the mesh resolution (nx x ny x nz) at 10x10x10µm, which is equivalent to half the average diameter of cardiomyocytes (Figure 2G).

The regularization parameter, E_reg_, was calibrated using an empirically determined value of λ = 0.01, and the tolerance and maximum number of iterations were set at 10^−8^ and 100, respectively, as stopping conditions. We used a four-level hierarchical approach as optimization method.

#### Optimization Strategy

To evaluate performance with resampled images, we used the same set of parameters for the MIRT algorithm, except for the grid resolution.

The grid resolution was set as the number of voxel equal to the average size of half a cardiomyocyte. Specifically, the images that were resampled at 10% had a grid resolution of 2-voxel, those at 15% and 20% had a grid resolution of 3-voxel, and the images resampled at 25% had a grid resolution of 4-voxel.

#### Image segmentation

We accurately segmented the cardiac tissue at frame (N/2), with N equal to the total number of time points in the image. Segmentation was performed manually using the open-source software ITK-SNAP (version 3.8.0) (Yushkevich et al., 2006) by the polygon inspector tool. We used 3D I/O ([IJ Plugins](https://ij-plugins.sourceforge.net/index.html)) in Fiji to convert the TIFF format in MetaImage Medical Format (.mha) format compatible with ITK-SNAP.

Segmentation was saved as The Neuroimaging Informatics Technology Initiative (.NIfTI) file (using the [nifti_io.jar](https://imagej.net/ij/plugins/nifti.html) plugin from Fiji) and imported into MATLAB. In MATLAB, the *fillholes3d* function was applied to the segmented images to fill any hole present by setting the maximum gap to be equal to 1 voxel.

#### Mesh Generation

To create a discretized version of heart tissue, we generated triangular (2-simplex element) meshes of each segmented heart using Iso2Mesh (Fang & Boas, 2009) a MATLAB/Octavebased mesh generation toolbox. Because myocardial tissue is sufficiently thin (30 µm or 1–2 cell layers), we created a surface mesh. We used a CGAL surface masher to create a genus-0 surface from the segmented image. The *vol2mesh* function was run with an isovalue fixed to 1, the maximum tetrahedral volume set to 4 voxels, and the maximum radius of the Delaunay sphere set to 4 voxels. Each surface was then smoothed with a Laplacian filter (*smoothsurf* function in Iso2Mesh), setting the smoothing parameter equal to 0.9. The algorithm was iterated 5 times. This step allowed us to have a smoother mesh surface and to eliminate possible inaccuracies associated with manual segmentation. The resulting mesh was then saved as Polygon File Format (.PLY) (*write_ply* function in toolbox_graph toolbox) and visualized in MeshLab (version 2020.06) (Cignoni et al., 2008), an open-source system for processing and editing 3D triangular meshes. In MeshLab, we run the Isotropic Explicit Remeshing function to improve aspect ratio (triangle quality) and topological regularity.

#### Selecting Myocardial Landmarks

We identified morphological points on the myocardial Live-Shape as follows:

- pt_1_-pt_2_ were taken at the border between the ventricle and the right and left IFT respectively;
- pt_3_ - pt_4_ were selected at the maximum elongation of the ventricle;
- pt_5_-pt_6_ were taken at the border between the cardiac mesoderm and the splanchnic mesoderm at the bending point of the IFT related to the foregut invagination;
- pt7 is a specific point located on the ventricle of the heart. This point was determined by the intersection of the myocardium and the longitudinal axis of symmetry, which was determined using Principal component analysis (PCA) in the mesh nodes. In cases where the shape of the heart was strongly asymmetric, due to missing parts of the IFTs, the point of asymmetry was manually corrected. We forced the shapes of the hearts to the centre of the axes, subtracting the mean values of the entire node’s distribution from each node. PCA defines the main 3 axis of the distribution of the nodes, considering the area of the mesh. The reason for considering the mesh area was that the faces of the mesh do not have the same area, particularly in smaller anatomic areas. Without weighing, PCA could imbalance the orientation of the heart shape, leading to inaccuracies in the determination of pt_7_.

#### Modelling the Staging System

We modelled the staging system as a Gaussian Mixture Model (GMMs). We examined whether the h/w values for each class followed a normal distribution. Due to the small number of specimens in each Atlas group (as reported in Table S3), we performed a Shapiro-Wilk test to determine normality with IBM SPSS Statistic (Shapiro & Wilk, 1965). The results showed no evidence of nonnormality, as shown in Table S4. Therefore, after a visual examination of Normal Q-Q plot, we decided to model the Atlas groups as a convex combination of normal distributions X_c_, parameterised by {µ_c_,σ_c_ ^2^} with c = 1…9 groups.

We associated the frames with the highest probability to the Atlas group. For the first and last frames where the matching is uncertain, we decided to classify only those with a probability of affiliation greater than 75%. We chose to exclude e06 due to its early stage of development. The e26 was excluded from the deformation analysis due to significant missing parts (IFTs). The staging of e01 was initiated from frame 11, as initial frames indicate that the embryos exhibit obsolete movements associated with stabilization outside the uterus. Conversely, e05 staging began at frame 14, since prior frames correspond to the period when the cardiac crescent is forming. During this period, the main motion estimation errors arise due to the incorporation of secondary heart field cells.

#### TMM Setting Parameters

The number of mixture components was determined by dividing the number of Live-Shape nodes in half. This number varied for each specimen and was determined by testing and adjusting to achieve a proper balance between computational efficiency and precision of registration. The EM algorithm was limited to a maximum of 150 iterations as a registration stop condition. The shapes were cantered on (0,0,0) before being registered, decreasing the computational cost.

#### Removing Missed Correspondences

After aligning the Atlas and Live-Shape using rigid registration, the shapes were imported into MeshLab and Atlas nodes that lacked correspondences with the Live-Shape were meticulously selected with the *Interactive Selection tool* and removed. The modified Atlas shape was referred to as Atlas_Cut_.

#### Defining the Live-Shape Mask

We applied the set of transformation fields {T_i_} to the segmentation of the myocardium relative to frame N/2 by a sequential B-Spline interpolations (with *mirt3D_transform* function in MIRT toolbox (Myronenko & Song, 2010)). This approach returned a 3D+t binary image of the myocardium during its shaping. We selected only the frames staged into the Atlas.

#### Defining the Atlas Mask

The Atlas binary mask was obtained by converting the Atlas_Cut_ shape into a 3D image. To achieve this, the Atlas_Cut_ was first restored to a genus-0 surface running MeshLab’s *closeholes* function. If the mesh had improper meshes, the shell had discontinuities that were manually closed. The mesh was then transformed to a shell of voxel in a 3D image using the *surf2volz* function in the Iso2mesh toolbox (Fang & Boas, 2009) with grid resolution equal to 1 voxel. The surface mask was then converted into a full binary mask by filling the gaps.

#### Principles of Finite Deformation Continuum Mechanics

To compute the deformation, we defined the rest triangular frame (R) and the deformed triangular frame (T) into the world space as:

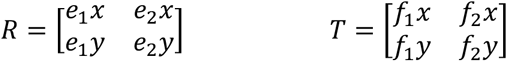

with e_1_ = r_2_-r_1_, e_2_ = r_3_-r_1_ with r_n_ equal to the 3D position of the element nodes. f_1_ = m_2_-m_1_, f_2_ = m_3_-m_1_, with m_n_ equal to the 3D position of the element nodes.

Then, we represented the deformation gradient F as the transformation from the rest triangular frame (R) into the deformed triangular frame (T):

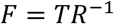

In particular, the growth rate J of a mesh element was determined by the Jacobian determinant of F. This value indicated how the area of the mesh element had changed due to the deformation. If the area is preserved, the deformation is isochoric, and J=1. On the other hand, J> 1 indicates tissue expansion, while J< 1 indicates tissue compression. Last, we computed the strain tensor ε. A classic way of measuring strain, limited to a little amount of deformation, is with the right Cauchy-Green strain tensor defined as:

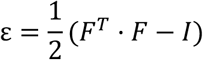

where I is the undeformed triangular frame (identity matrix). ε is a matrix that provides information about the stretching and shearing of each mesh element. Looking at the diagonal values, we defined the main stretches and their direction as eigenvalues and eigenvectors for each mesh element. The anisotropy was determined by the ratio of its two main eigenvalues, which was denoted as θ. θ = 1 indicates isotropic deformation. θ > 1 indicates anisotropic deformation.

All these deformation measurements were translation and rotation invariant; therefore, they were not affected by relative movements of the embryo during image acquisition.

To minimize distortions related to isolated events, such as uncoordinated beating of cells during morphogenesis, we smoothed the deformation values within a geodesic distance up to 20 voxels.

The deformation pattern was calculated between two time-points and always mapped onto the shape of the last time considered.

#### Concatenating Individual Motion Profiles to Compute the Cumulative Deformation

To identify anatomical correspondences in the Atlas across the groups, we used motion profiles from different live images. SurfaceMaps represented individual motion profiles projected onto the Atlas. We established Atlas correspondences by aligning multiples SurfaceMaps. Equally staged SurfaceMaps had the same geometry, but different orientation and point density. To address this variability, we designated one SurfaceMap as the reference, known as eye-PC, and concatenated it with the hook-PC in a three-step process (Figure 4B).

1) We used the [TMM-based rigid registration](https://es.mathworks.com/matlabcentral/fileexchange/63693-robust-group-wise-registration-of-point-sets-using-multi-resolution-t-mixture-model?s_tid=prof_contriblnk) to align each hook-PC with the eye-PC (Ravikumar et al., 2016, 2018). This enabled us to overcome the differences in orientation between the two point-clouds; 2) We converted the registered hook-PC into binary masks and used a non-rigid registration algorithm to extract the T-transformation needed to morph each mask into the e27 SurfaceMaps mask (Myronenko et al., 2007); 3) We employed a k-nearest neighbours algorithm (*knnsearch* function in MATLAB) to establish a face-to-face match between each hook-PC and the eye-PC. This step was crucial in addressing the differences in the density of points between the two point-clouds.

Referring to Figure 4C, we started the concatenation path at Gr_4_ of embryo 27 (e27), using its SurfaceMap as our spatial reference. Embryos e02, e16, e124, e22, and e35 each had a staged frame in Gr_4_. Through a face-to-face spatial mapping process, we projected the spatial reference (the e27 SurfaceMap) onto the SurfaceMaps of these other embryos (e02, e16, e124, e22, and e35) by identifying anatomical correspondences.

These correspondences between the eye-PC (e27) and the hook-PC (e02, e16, e124, e22, and e35) allowed us to indirectly determine the position of the e27 SurfaceMap in Gr_3_.

To compare the deformation patterns of e31 and e05 in Gr_3_, we used the anatomical correspondences in the e22 SurfaceMap as the eye-PC and applied face-to-face mapping between the SurfaceMaps of e31 and e05 and the eye-PC.

We used the same method to estimate the distribution of e27 SurfaceMap in Gr_6_, this time using the e24 SurfaceMap for face-to-face mapping. To calculate the distribution in Gr_8_, we fixed the anatomical correspondences in the e24 SurfaceMap as the eye-PC.

#### Concatenating Individual Motion Profiles to Compute in-Silico Fate Map

To concatenate the different specimens and create a pseudo-cell tracking, we used their SurfaceMaps. As previously discussed, while the SurfaceMaps within the same group had similar shapes, they had different density of points.

To address this issue and establish correspondences between SurfaceMaps, we designated the SurfaceMaps of one embryo, e35, as the reference point cloud.

This decision was based on the fact that the live images of e35 covered a central sub-window of the developmental timeline. Therefore, we used e35 SurfaceMaps as the distribution of points to describe the motion profile of Atlas.

We followed the same concatenation approach as before, defining the eye-PC and the hook-PC. We started by matching the SurfaceMap of e35 with that of e16 at Gr_4_. Next, we used the motion profile of e16 to infer the position of the SurfaceMap of e35 at Gr_3_. This became the new eye-PC to which the SurfaceMaps of e31 were anchored, and so on.

By combining the individual motion profiles, we created a dynamic Atlas. The dynamic Atlas maintained the original morphology of the static Atlas, but the positions of the mesh nodes were altered following the motion profile derived from the concatenation of the motion profiles of the live embryos.

### Computational Resources

The analyses were performed on a workstation equipped with an Intel(R) Xeon(R) E-2176M CPU operating at 2.70 GHz (2712 MHz), featuring 6 physical cores and 12 logical processors, 64 GB of RAM, and a NVIDIA Quadro P4200 GPU.

## Acknowledgements

We thank members of the Torres group for inspiring discussions and advice. We thank Peter Majer (Bitplane) for helpful advice and guidance on the work performed. We thank members of the Microscopy and Dynamic Imaging, Transgenesis, and Animal Facility CNIC units for excellent support. M.R. was a recipient of a Marie Skłodowska-Curie postdoctoral contract from H2020-MSCA-ITN-2016-722427. This work was funded by grants PGC2018-096486-B-I00 and PID2022-140058NB-C31 from the Agencia Estatal de Investigación to M.T.; Comunidad de Madrid grant P2022/BMD-7245 CARDIOBOOST-CM to M.T.; European Research Council AdG ref. 101142005 to M. T. The CNIC Unit of Microscopy and Dynamic Imaging is supported by FEDER ‘Una manera de hacer Europa’ (ReDIB ICTS infrastructure TRIMA@CNIC, MCIN). The CNIC is supported by the Instituto de Salud Carlos III (ISCIII), the Ministerio de Ciencia, Innovación y Universidades (MICIU) and the Pro CNIC Foundation, and is a Severo Ochoa Center of Excellence (grant CEX2020-001041-S funded by MICIU/AEI/10.13039/501100011033).

## Author contributions

Conceptualization: M.R., M.T.; Methodology: M.S., M.R.; Software: M.R.; Formal analysis: M.R.; Investigation: M.S., I.E., K.I., M.T.; Data curation: M.R., M.T.; Writing - original draft: M.R.; Writing - review & editing: M.R., M.T.; Supervision: M.T.; Project administration: M.T.; Funding acquisition: M.T.

## SUPPLEMENTARY FIGURES

**Figure S1.**
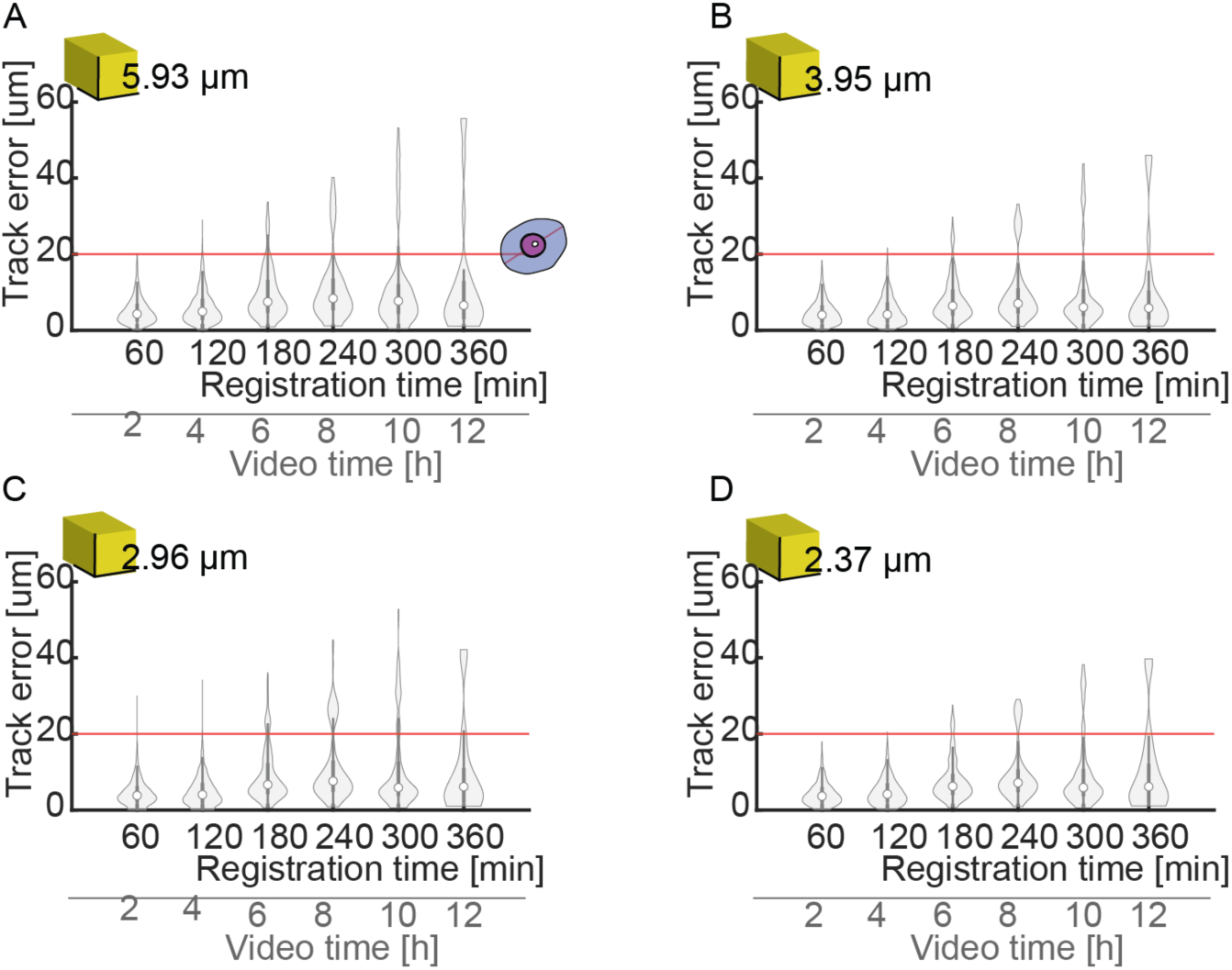
Error estimation of motion profile computed for the resampled images. The sequential error was estimated for the ground-truth points when the image was resized at 10%(A), 15%(B), 20%(C), and 25%(D). The error is expressed in μm. The red line indicates the averaged cell diameter of 20 μm. The yellow cubes indicate the voxel size. The registration time is half of the video time, according to the registration strategy adopted. The graphs represent the errors of 10086 tracked cells in the images of 9 embryos (e01, e02, e05, e06, e15, e16, e24, e26, e27).

**Figure S2.**
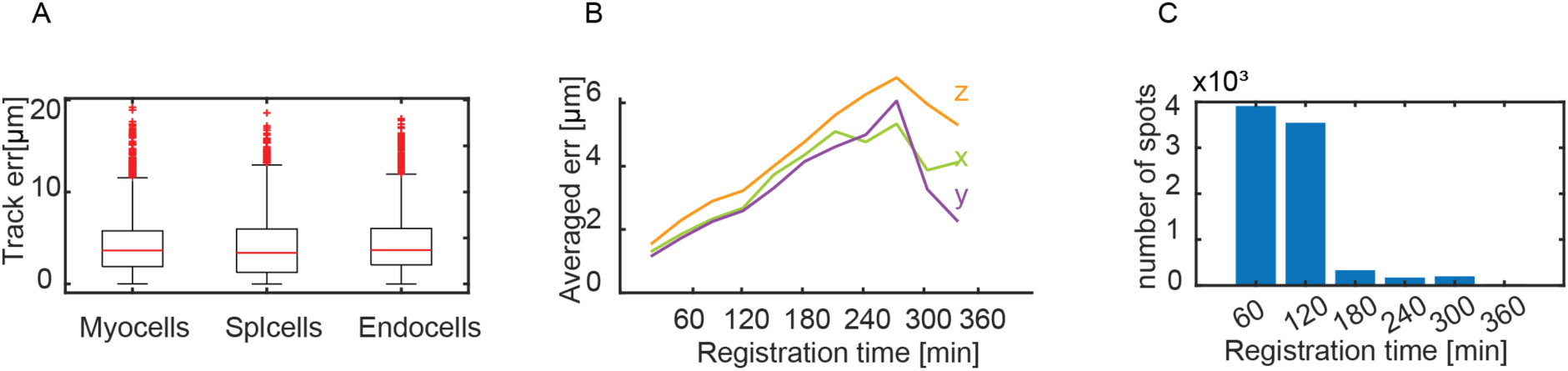
Motion error estimation. (A), The error made in continuous registration of landmarks belonging to different cardiac layers from images at 25% resolution. There is no statistically significant difference between the error made on myocardial cells, splanchnic mesodermal cells, and endodermal cells. (B), x-y-z decomposition of the continuous mean error, related to a 25% rescaled image (e01, e02, e05, e06, e15, e16, e24, e26, e27) is reported. (C) Number of landmarks in two-hours video interval for which the error was estimated.

**Figure S3.**
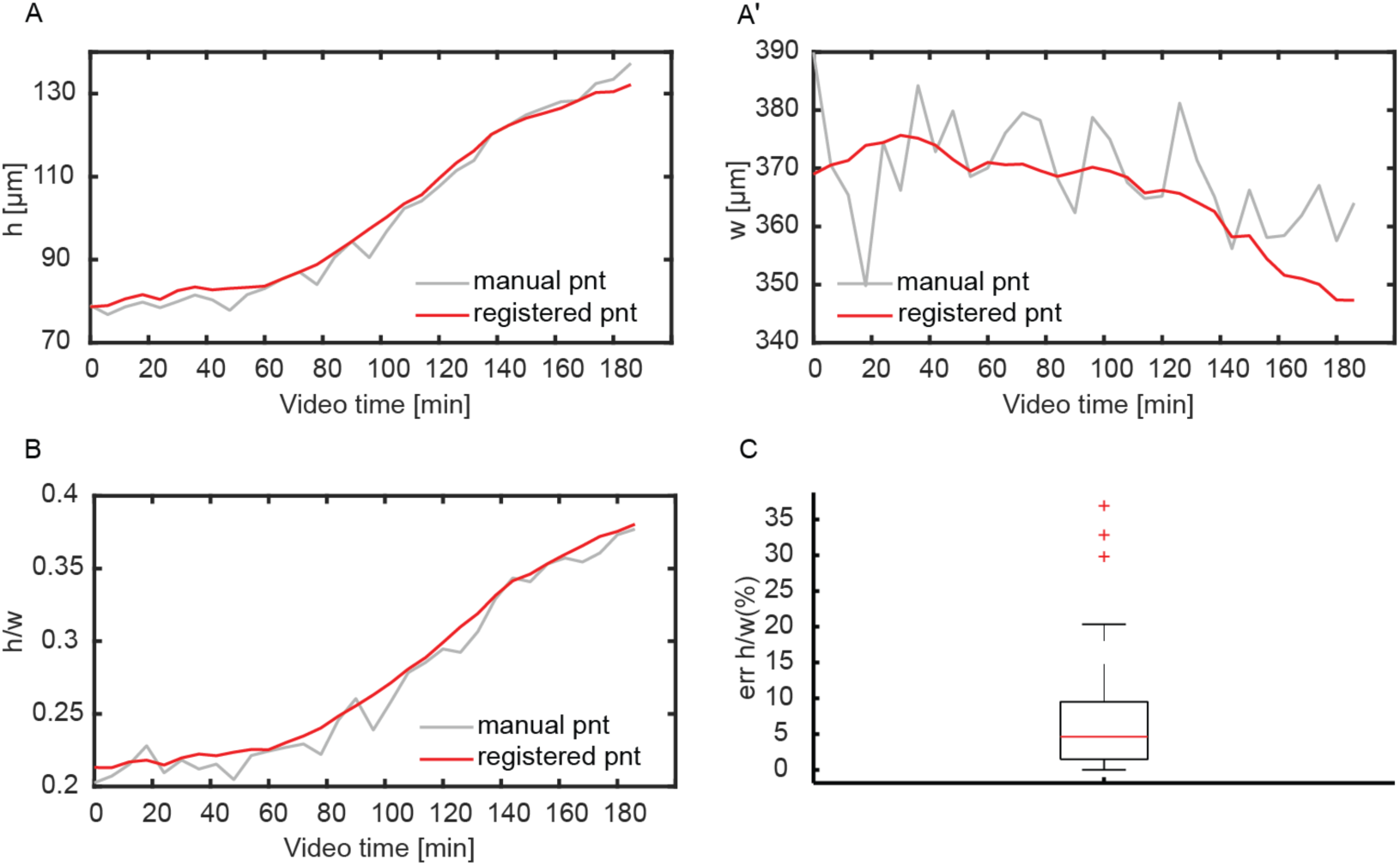
Evaluation of h/w feature selected manually or computed by the motion profile. The data are related to a single embryo (e01). (A,A’), In red the h and w lengths between landmarks taken manually on the shapes at each frame. In grey, the lengths h and w calculated automatically for each frame at the selected landmark at time t0 (registered point). (B), Comparison between the feature h/w obtained from the manual landmarks (red line) and the landmarks defined by the continuous model (gray line). (C), Percentage difference between the two features performed on 3 embryos (e01, e07, e27).

**Figure S4.**
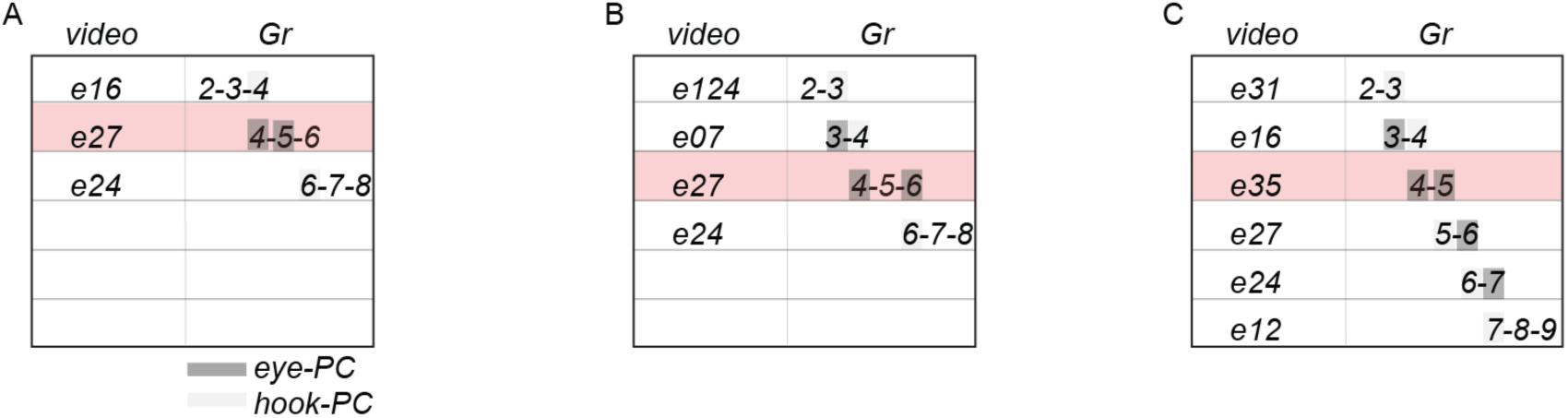
Concatenation paths defining fate map. We evaluated the growth profile for the 10 zones between Gr2 to Gr8 for three different concatenation paths, shown in (A), (B), (C). We compare Gr2 to Gr8 as it represents the common stage range among the three different paths. The matching score between the growth profiles of the model and the one extracted from the live image was 62%, 65% and 81%, respectively. We chose the last concatenation path because it showed the highest match.

**Figure S5.**
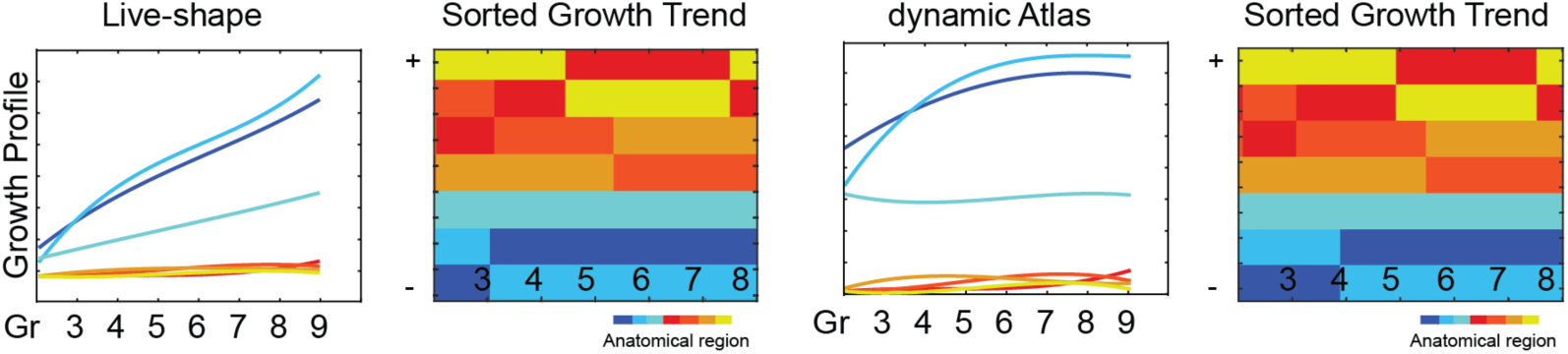
Validating the Fate Map at Regional Level. The growth profile for the anatomical regions (excluding regions 1,5, and 10) were computed for the live-shapes and the Dynamic Atlas, with the corresponding heatmap. The two sorted growth trend matches at 92%.

## SUPPLEMENTARY TABLES

**Table S1.**
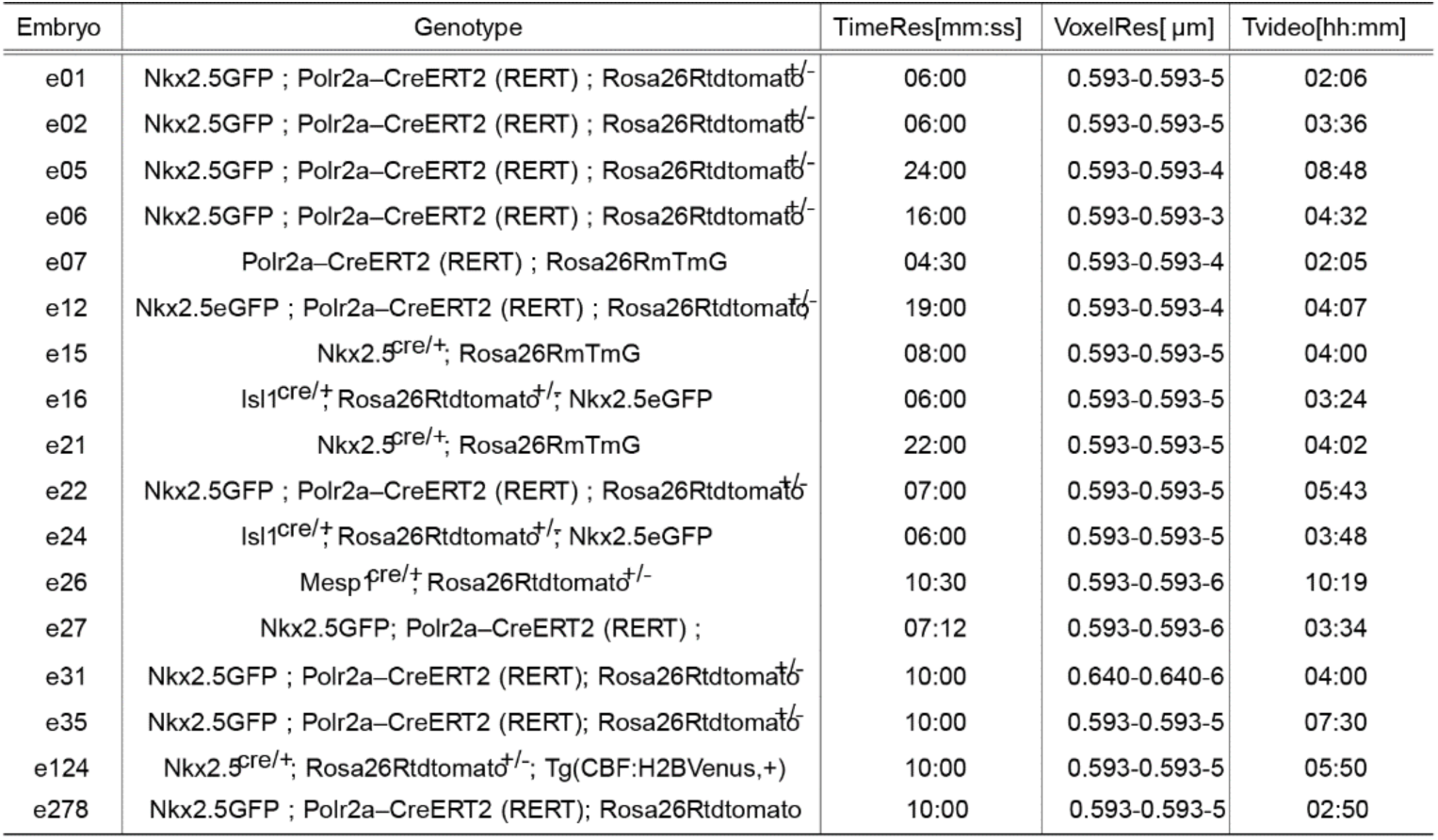
Dataset of live images. . The first column shows the list of embryos involved in our study; Genotype column reports the mouse allels (Nkx2.5GFP(Wu et al., 2006), Polr2a-CreERT2 (RERT)(Guerra et al., 2003), Nkx2.5cre/+(Stanley et al., 2004), Rosa26Rtdtomato+/-(Madisen et al., 2010), Rosa26RmTmG(Muzumdar et al., 2007), Tg(CBF:H2BVenus,+)(Nowotschin et al., 2013)); TimeRes column lists the time resolution of the time-lapse[mm:ss]; VoxelRes column indicates the resolution in pm along the x,y,z axes; the last column (T_video_) shows the total duration of the time-lapse[hh:mm]. e278 in grastrulation phase.

**Table S2.** Summary of tracked cells in different tissues. We have indicated the number of frames for each video (N_frames_), the duration of the video (T[hh:mml), the number of total tracked cells (NumCell). We have indicated which of these cells belong to the myocardium (Myo), how many to the mesoderm splachnic (Spl) and how many to the endoderm (Endo). In Totcells the total number of points, i.e.. cells, for which the error is quantified. The total is given by the number of cells Numcells*Nframes. For e05 and e15, 1 and 4 endothelial cells were tracked respectively.

| Em-bryo | Nframes | T[hh:mm] | Num-Cell | Myo | Spl | Endo | Tot-cells |
| --- | --- | --- | --- | --- | --- | --- | --- |
| e01 | 22 | 02:06 | 83 | 50 | 13 | 20 | 2656 |
| e02 | 37 | 03:36 | 30 | 7 | 10 | 20 | 1110 |
| e05 | 23 | 08:48 | 10 | 8 | 1 | - | 230 |
| e06 | 18 | 04:32 | 30 | 8 | - | 22 | 540 |
| e15 | 31 | 04:00 | 12 | - | - | - | 372 |
| e16 | 35 | 03:24 | 56 | 22 | 11 | 25 | 1960 |
| e26 | 59 | 10:19 | 11 | - | - | - | 616 |
| e24 | 39 | 03:48 | 30 | 23 | 2 | 12 | 1170 |
| e27 | 31 | 03:34 | 30 | 23 | 5 | 1 | 930 |

**Table S3.** ATLAS staging system results. Number of specimens used to build the ATLAS Grs (Esteban et al., 2022).

| Gr | NumSpecimen |
| --- | --- |
| 1 | 5 |
| 2 | 5 |
| 3 | 4 |
| 4 | 5 |
| 5 | 5 |
| 6 | 7 |
| 7 | 5 |
| 8 | 5 |
| 9 | 7 |
| 10 | 2 |

**Table S4.**
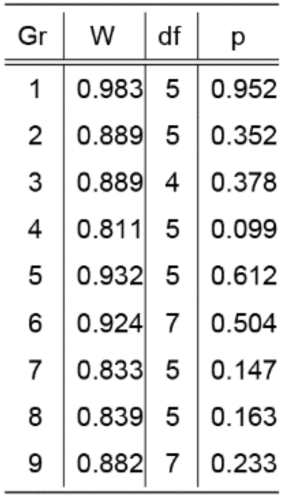
Shapiro-Wilk test results for each ATLAS group.

| Gr | W | df | p |
| --- | --- | --- | --- |
| 1 | 0.983 | 5 | 0.952 |
| 2 | 0.889 | 5 | 0.352 |
| 3 | 0.889 | 4 | 0.378 |
| 4 | 0.811 | 5 | 0.099 |
| 5 | 0.932 | 5 | 0.612 |
| 6 | 0.924 | 7 | 0.504 |
| 7 | 0.833 | 5 | 0.147 |
| 8 | 0.839 | 5 | 0.163 |
| 9 | 0.882 | 7 | 0.233 |

**Table S5.** Comparison of Point Cloud rigid registration algorithms. In each column, the rmse is reported for each embryo at each registration technique (ICP, CPD, TMM). We assessed the performance of the most popular methods for rigid point set registration: ICP(Segal et al., 2009) and the CPD(Myronenko & Song, 2010b) and TMM-based(Ravikumar et al., 2018) algorithms. To this end, we evaluated how well the algorithms aligned a shape to a modified version of that shape. The altered shape underwent three modifications: 1) a random reduction of 30% of its points. 2) a rigid transformation as a rotation of 20° around zaxis; rotation of 70° around the x-axis; a rescale of 20% with respect to the original shape size. 3) a partial cut of the IFTs. The algorithms were tested to the first frame (t0) of five different embryos (e01, e02, e15, e16, e27) by calculating the root-mean-square error (rmse) between the original shape and the corresponding points in the altered one We found that the TMM-based algorithm performed better compared to ICP and CPD algorithms. So, we decided to implement the TMM-based algorithm as the first step in the spatial mapping pipeline.

|  | Rotating + Resizing |  |  | Rotating + Resizing + Cutting |  |  |
| --- | --- | --- | --- | --- | --- | --- |
|  | ICP | CPD | TMM | ICP | CPD | TMM |
| e01 | 5.130826 | 9.747474 | 2.263551 | 24.12582 | 8.197942 | 1.603977 |
| e02 | 14.88792 | 10.45908 | 2.979164 | 15.09942 | 8.55301 | 2.922606 |
| e15 | 14.50765 | 12.64864 | 1.873373 | 10.09741 | 11.42981 | 1.396486 |
| e16 | 4.04974 | 8.714035 | 2.174541 | 10.89691 | 7.318639 | 1.413487 |
| e27 | 5.532653 | 11.20625 | 1.640565 | 25.85086 | 9.455913 | 1.241526 |

**Table S6.** Staging system result. The first column shows the ATLAS group(Gr) associated with the live embryo shown in the second column. The last two columns show the video frames related to the rest state (frame(Gr-1)) and the deformed state (frame(Gr)), respectively.

| Gr | embryo | frame(Gr-1) | frame(Gr) |
| --- | --- | --- | --- |
| 2 | e21 | 1 | 11 |
| 2 | e22 | 3 | 34 |
| 3 | e05 | 19 | 22 |
| 3 | e22 | 34 | 40 |
| 3 | e31 | 18 | 23 |
| 3 | e16 | 1 | 12 |
| 3 | e124 | 14 | 23 |
| 4 | e02 | 9 | 26 |
| 4 | e16 | 12 | 27 |
| 4 | e22 | 40 | 46 |
| 4 | e35 | 6 | 20 |
| 4 | e124 | 23 | 32 |
| 5 | e07 | 9 | 21 |
| 5 | e27 | 7 | 19 |
| 5 | e35 | 20 | 33 |
| 6 | e27 | 19 | 29 |
| 7 | e24 | 22 | 34 |
| 8 | e12 | 3 | 7 |
| 8 | e24 | 34 | 39 |
| 9 | e12 | 7 | 13 |

**Table S7.** Validating the fate map at cellular level.

| Embryo | NumTrackedCells | RightPredicted |
| --- | --- | --- |
| e01 | 13 | 12 |
| e02 | 7 | 7 |
| e05 | 6 | 7 |
| e16 | 13 | 11 |

## SUPPLEMENTARY VIDEOS

Video S1: **HT Morphogenesis as a Continuum**. On the right, the live image of e01 captures the dynamic tissue movements during morphogenesis. On the left, the discretized model of the HT illustrates the motion profile.

Video S2: **Spatio-temporal referencing of live images into the Atlas**. Frames 7, 19, and 29 of embryo e27 (grey shapes and grey mesh) are staged in Gr_4_, Gr_5_, and Gr_6_ of the Atlas. The staged frames are then mapped face-to-face into the Atlas (blue shapes). A yellow spot marks the corresponding anatomical position across the live images, live shapes, and the Atlas.

Video S3: ***In-silico* Fate Map. The dynamic Atlas is represented in its point cloud version**. At the cellular level, a yellow spot tracks a hypothetical cell position throughout heart morphogenesis. At the tissue level, yellow spots illustrate the tissue deformation occurring during heart morphogenesis.

